# Velocity-dependent friction in a simple two-oscillator system. I. Phase–amplitude coupling

**DOI:** 10.64898/2026.11.18.767449

**Authors:** Yuhan Luo, Sonali Priyadarshini Nayak, Murielle Hsu, Gregory A. Worrell, David Hsu

## Abstract

Cortical gamma oscillations often arrive in bursts locked to the phase of slower theta waves, a pattern called theta–gamma phase–amplitude coupling (PAC). PAC is linked to memory and altered in epilepsy, but most models tie it to particular cells or circuits. We start from a generalized Langevin equation, an equation of motion for a few collective variables in which friction and random forces stand for the unobserved neural activity, and let its stiffness and friction depend on the state. Expanding them to first order in the theta and gamma variables yields ten possible interaction terms; only the two friction-type terms change gamma energy and produce PAC, and depending on the term and its sign the gamma bursts sit at the peak, trough, rising or falling phase of theta. We study a gamma friction that follows theta velocity as a simplified example. Because the time integral of theta velocity is theta coordinate, the gamma amplitude rises and falls with theta coordinate as exp(*g*_*vv*_*x*_*θ*_*/*2). The fluctuation–dissipation theorem turns noise and damping into a theta effective temperature, and the PAC strength becomes a closed-form function of one dimensionless number, *κ*, that combines the coupling with this temperature. Without fitted parameters, this prediction accounts for 51,095 simulated records (*R*^2^ = 0.993) and for the median PAC of 5,160 combinations of coupling and damping (*R*^2^ = 0.989). The coupling also drives gamma out of equilibrium: its kinetic temperature grows far above the one inferred from its response. All coefficients enter linearly, so that they can in principle be estimated from recordings by linear regression. A companion paper shows that the same number sets the distance to a stability boundary.

## 1. Introduction

Phase–amplitude coupling (PAC) has been described in the brain across multiple frequencies and brain regions [1–4]. Of these, the coupling of the amplitude of gamma oscillations (30 to 100 Hz) to the phase of theta oscillations (4 to 8 Hz) in the hippocampus is the most prominent and best studied, and is thought to play a role in information encoding, sequencing and retrieval [5–11]. Theta–gamma coupling is altered in ictogenesis in a complicated way. For instance, theta–gamma coupling increases early in a seizure [12, 13], and externally applied electrical stimulation at theta frequencies can provoke seizures in an epileptic brain [14]. On the other hand, theta–gamma coupling is impaired in a mouse model of Dravet syndrome [15], and a well-defined hippocampal theta band is severely degraded with epileptogenesis in a kainic acid rat model [16].

Mathematical models for theta–gamma coupling include neuronal and neural mass models [17–22] as well as phenomenological nonlinear models, particularly notable among the latter being the Epileptor model [23], the model parameters of which can be personalized for individual patients through Bayesian statistics [24, 25]. Macroscopic models such as the neural mass and Epileptor models have the advantage of being simpler to analyse, with the hope that some system-wide properties may be more apparent at the macroscopic level than at the microscopic level [26, 27].Macroscopic and microscopic models provide complementary descriptions of a system across different length and time scales. Minimal dynamical models have been employed to investigate emergent behavior in complex systems, ranging from cross-frequency coupling in neuronal oscillations[28] to regulatory dynamics and transitions between distinct states in diverse complex systems[29–31].

Here, we investigate a simple macroscopic mathematical model, that of two damped oscillators, one oscillating at theta frequency (4 Hz), the other at gamma frequency (60 Hz), both driven by Gaussian random noise. When the interaction between the two oscillators is linear in the coordinates of the oscillators, the interaction shifts the oscillator frequencies but does not produce true PAC. Here we generalize the interaction between the two oscillators to include the lowest order nonlinear interactions, and ask if any of these terms lead to strong PAC. If the answer is yes, then this simple model may be further generalized to a wider family of generalized Langevin equations (GLE) [32, 33], and one may hope to build a GLE model for brain dynamics. Within a GLE framework, for example, friction may represent inhibitory interneuronal interactions. When analyzing data from electroencephalograms (EEG), the GLE coordinate represents an electric potential measured by EEG electrodes, while the velocity represents the time rate of change of the electric potential.

## 2. Theory: a second-order model of theta and gamma oscillators

### 2.1. The generalized Langevin equation and its expansion

The generalized Langevin equation can be written as

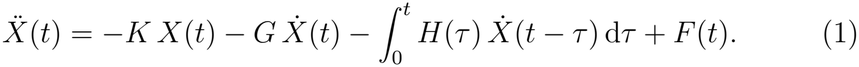

In this form, *X*(*t*) is a time-dependent vector comprised of the chosen experimental observables, here the theta and gamma components of a recorded EEG signal, and *K, G, H* and *F* are empirical parameters that depend on the specifics of a particular neural system. *K* plays the role of a linear restoring force, giving rise to oscillations; *G* plays the role of a friction, which slows the dynamics; *H* is a memory function, which allows the past to affect the future; and *F* (*t*) is a random force, representing a thermal bath [32, 33]. Because these parameters enter the equation of motion linearly, they can be extracted from experimental data by linear regression [Hsu, Techarxiv manuscript].

As it stands, Eq. (1) is linear in *X*. Cross-oscillator coupling produces a new set of oscillators but not PAC. To explore the effect of nonlinear interactions, we expand *K, G* and *H* as functions of *X* and 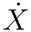:

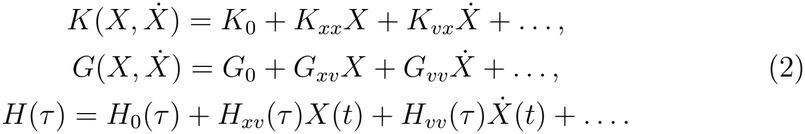

Each coefficient carries three indices: 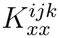, for example, multiplies *X*_*k*_*X*_*j*_ in the equation of motion for component *i*. A product of a coordinate and a velocity can be written in *K* or in *G*; we attribute it to *K*_*vx*_ when component *i* is a coordinate, and to *G*_*xv*_ when component *i* is a velocity.

### 2.2. Ten possible interactions of three kinds

For the rest of this paper, we will ignore the memory term. We do not imply that memory is unimportant in the brain (memory is certainly important in the brain), but we wish first to understand the effect of adding in just the lowest order nonlinear terms, which are those that are bilinear in the system coordinate and/or velocity, before we consider complexities introduced by memory.

For a theta oscillator and a gamma oscillator, the equation of motion for the gamma oscillator contains ten second-order terms. To see what each term does, we ask how it changes the gamma oscillation. Because the gamma oscillator is fifteen times faster than the theta oscillator, theta hardly changes within one gamma cycle, so each term can be judged by its average over one gamma cycle with theta held fixed (Supplementary Section S2). The terms fall into three kinds.

i. *Friction type, G*_*vv*_*v*_*θ*_*v*_*γ*_ and *G*_*xv*_*x*_*θ*_*v*_*γ*_. These terms are frictional terms, proportional to the gamma velocity. If *G*_*vv*_*v*_*θ*_ or *G*_*xv*_*x*_*θ*_ is positive, then the friction acting on the gamma oscillator is increased above baseline (*G*_0_); if negative, then the friction acting on the gamma oscillator is decreased below baseline. When the friction is above baseline, the gamma oscillator will be suppressed, and its amplitude of oscillation will decrease. When the friction is below baseline, the gamma oscillator will oscillate at higher amplitude. Since both *G*_*vv*_*v*_*θ*_ and *G*_*xv*_*x*_*θ*_ change sign with the phase of the theta oscillation, these two friction terms introduce theta–gamma PAC.
ii. *Stiffness type, K*_*xx*_*x*_*θ*_*x*_*γ*_ and *K*_*vx*_*v*_*θ*_*x*_*γ*_. These terms change the effective spring constant of the gamma oscillator, thus producing phase–frequency coupling, but not PAC.
iii. *Single-oscillator terms*, 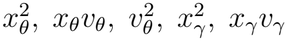, and 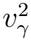. These terms produce no cross-oscillator coupling, and will not produce PAC.

Here and below, *G*_*vv*_, *G*_*xv*_, *K*_*xx*_ and *K*_*vx*_ denote the coefficients of the gamma equation of motion.

### 2.3. The simplified model

Now let us focus on the velocity–velocity term *G*_*vv*_ to study its effect on PAC, and temporarily drop the other bilinear terms. The effect of *G*_*xv*_ on PAC will be similar, but shifted by a quarter phase. The equations of motion are then:

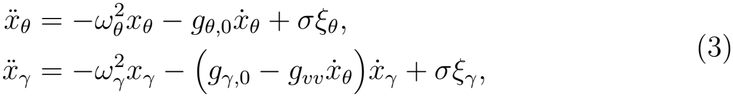

The theta and gamma oscillators are driven by independent Gaussian white noises *ξ*_*θ*_ and *ξ*_*γ*_, with angular frequencies *ω*_*θ*_ and *ω*_*γ*_, baseline frictions *g*_*θ*,0_ and *g*_*γ*,0_, and noise intensity *σ*. Note that *g*_*vv*_ = −*G*_*vv*_. In this model, the friction acting on the gamma oscillator depends on the velocity of the theta oscillator through an effective friction coefficient *g*_*γ*,eff_ = *g*_*γ*,0_ −*g*_*vv*_*v*_*θ*_. If *v*_*θ*_ is positive and *g*_*vv*_ is also positive, the coupling reduces the effective friction acting on the gamma oscillator, so that its amplitude grows; when *v*_*θ*_ is negative, the coupling increases the effective friction, so that the amplitude shrinks (Fig. 1). When *g*_*vv*_ is positive, the gamma oscillator thus gains energy while the theta oscillator rises and loses energy while it falls; gamma oscillations are thus amplified at theta peaks, and suppressed at theta troughs.

**Figure 1:**
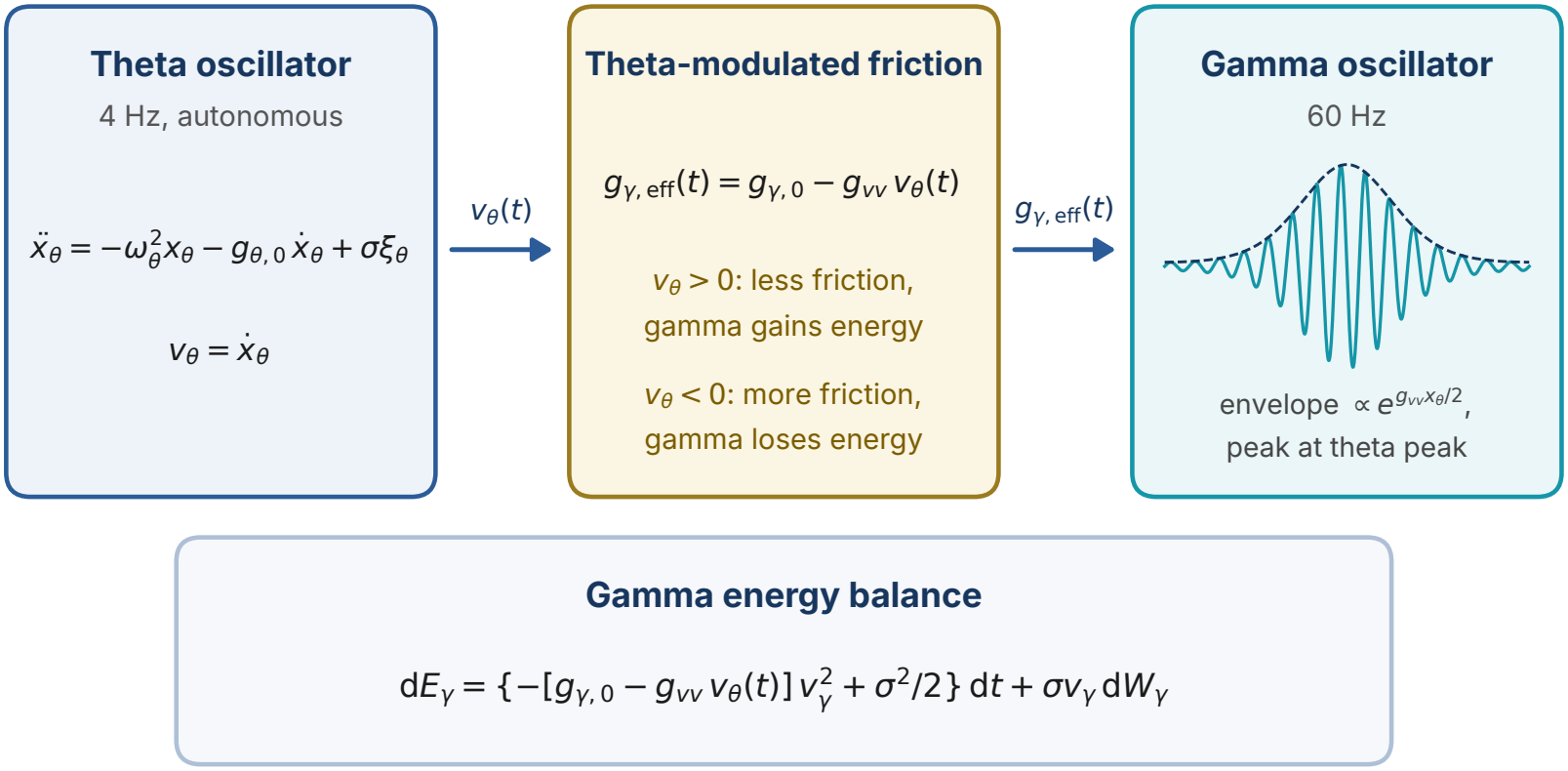
Framework of the model. Theta velocity modulates gamma friction within each theta cycle, so gamma gains energy while theta coordinate rises and loses it while theta coordinate falls; the gamma envelope therefore peaks at the theta coordinate peak.

For convenience, we define baseline friction in terms of a single damping ratio *ζ* in terms of which *g*_*θ*,0_ = *ζω*_*θ*_ and *g*_*γ*,0_ = *ζω*_*γ*_. Critical damping corresponds to *ζ* = 2, below which the oscillators are underdamped and above which they are overdamped. For brevity, we refer to the variables of the theta and gamma oscillators as theta and gamma variables (for example, theta velocity). Table 1 lists the quantities used in the rest of the paper.

**Table 1:**
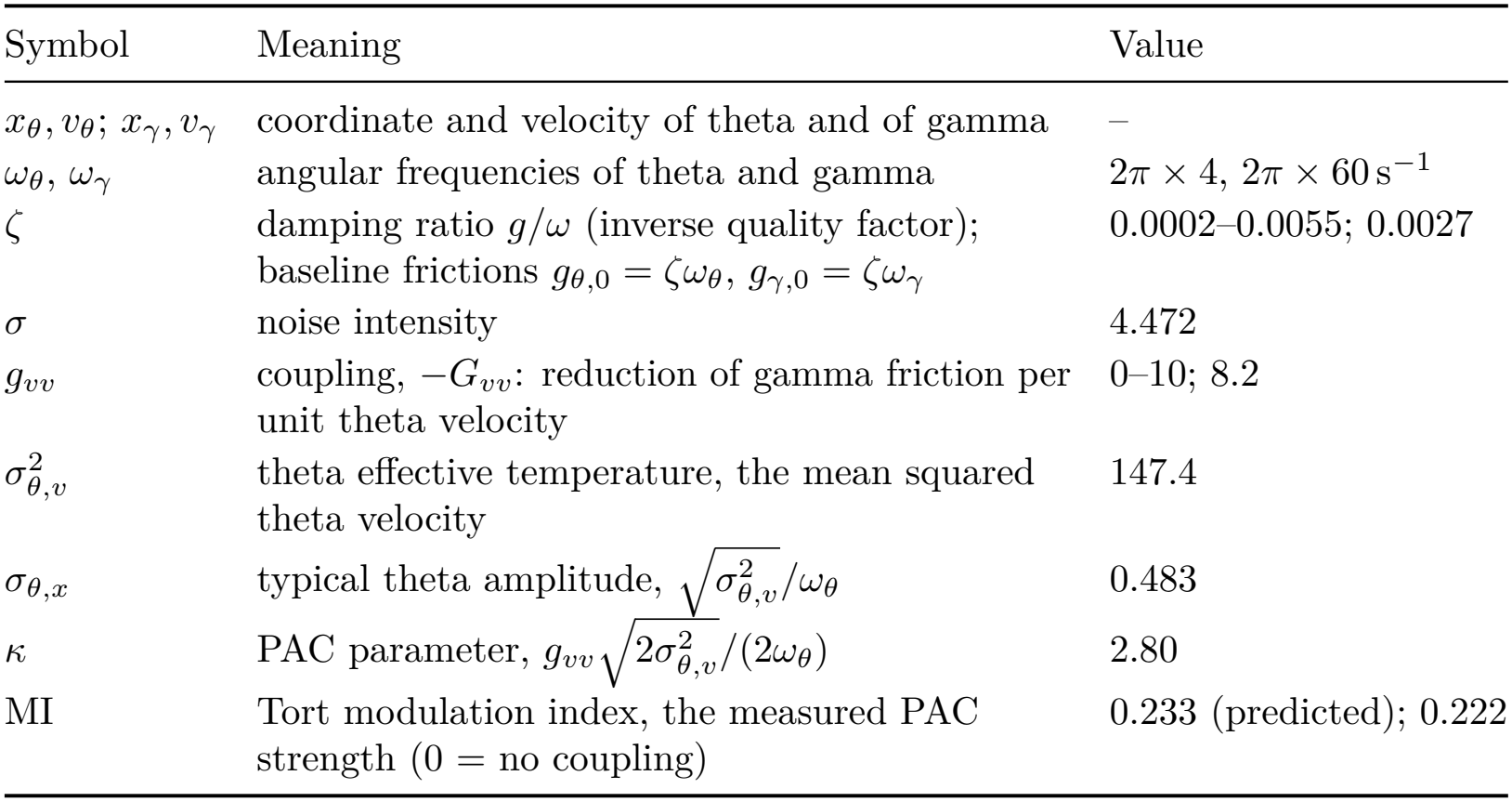
Main quantities. Values refer to the representative point C1 (*g*_*vv*_ = 8.2, *ζ* = 0.0027) unless stated otherwise.

| Symbol | Meaning | Value |
|---|---|---|
| *x*<sub>θ</sub>, *v*<sub>θ</sub>; *x*<sub>γ</sub>, *v*<sub>γ</sub> | coordinate and velocity of theta and of gamma | – |
| ω<sub>θ</sub>, ω<sub>γ</sub> | angular frequencies of theta and gamma | 2π × 4, 2π × 60 s<sup>−1</sup> |
| ζ | damping ratio *g*/ω (inverse quality factor);<br>baseline frictions *g*<sub>θ,0</sub> = ζω<sub>θ</sub>, *g*<sub>γ,0</sub> = ζω<sub>γ</sub> | 0.0002–0.0055; 0.0027 |
| σ | noise intensity | 4.472 |
| *g*<sub>*vv*</sub> | coupling, −*G*<sub>*vv*</sub>: reduction of gamma friction per unit theta velocity | 0–10; 8.2 |
| σ<sup>2</sup><sub>θ,*v*</sub> | theta effective temperature, the mean squared theta velocity | 147.4 |
| σ<sub>θ,*x*</sub> | typical theta amplitude, $\sqrt{\sigma_{\theta,v}^2}/\omega_\theta$ | 0.483 |
| κ | PAC parameter, $g_{vv}\sqrt{2\sigma_{\theta,v}^2}/(2\omega_\theta)$ | 2.80 |
| MI | Tort modulation index, the measured PAC strength (0 = no coupling) | 0.233 (predicted); 0.222 |

## 3. What the model predicts

This section derives three results, each from the one before: the size of theta fluctuations, the shape of the gamma envelope and the PAC strength. We give the core equations and their meaning, and the complete derivations are in Appendix A and Supplementary Section S3.

### 3.1. Theta behaves like a particle in a heat bath

The theta oscillator is driven by noise and damped by friction. At steady-state, the fluctuation-dissipation theorem (FDT) states that these two effects must be balanced, so that the energy gained through the random noise forcing term is on average equal to the energy dissipated through friction [32, 34]. This energy balance for the theta oscillator results in the following well-known expressions for the mean squared velocity and coordinate:

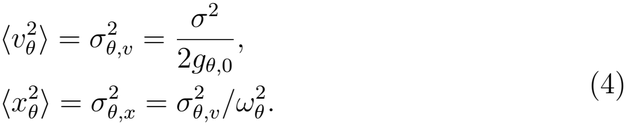

Note also that 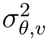 is equal to the temperature of the system. Increasing noise increases the temperature of the system while increasing friction reduces temperature.

### 3.2. The gamma amplitude envelope follows theta coordinate

Let *a*_*γ*_ be the amplitude envelope of the gamma oscillator, which is damped by an effective friction, defined as *g*_*γ*,eff_ = *g*_*γ*,0_ −*g*_*vv*_*v*_*θ*_ (see Eq. (3)). When *g*_*vv*_ *>* 0 and *v*_*θ*_ *>* 0, then the effective friction is reduced from its baseline value of *g*_*γ*,0_. When *v*_*θ*_ changes sign and becomes negative, then the effective friction is increased from its baseline value of *g*_*γ*,0_. This change in the effective friction over a full theta cycle gives rise to theta-gamma PAC. In Appendix A.2, we show that the part of the gamma amplitude envelope that gives rise to theta–gamma PAC is

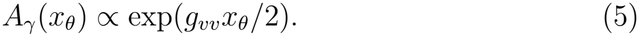

Eq. (5) is a central result of this paper.

### 3.3. One dimensionless number determines PAC strength

We quantify theta–gamma PAC by the Tort modulation index (MI) [9]: the theta cycle is divided into 18 phase bins, the mean gamma amplitude envelope is computed in each bin, and the Tort MI is then calculated, yielding a measure of the deviation of this phase amplitude profile from a flat one (MI = 0, no PAC), to perfect correlation (MI = 1). In Appendix A.4, we show that MI can be approximated as follows:

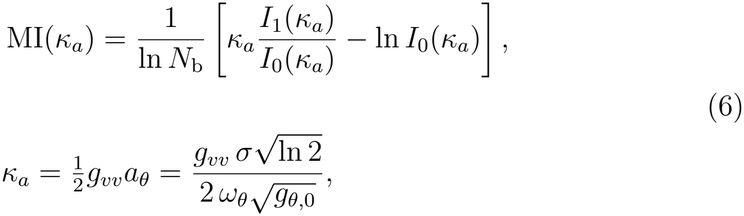

Here *N*_b_ = 18 is the number of theta-phase bins and *I*_0_, *I*_1_ are modified Bessel functions. The expression above for the dimensionless number *κ*_*a*_ is a second central result of this paper. Unsurprisingly, Eq. (6) predicts that MI increases with larger values of *g*_*vv*_ and smaller values of *ω*_*θ*_ and *g*_*θ*,0_. Somewhat surprisingly, Eq. (6) also predicts that MI should increase with increased noise intensity, because stronger noise makes the theta oscillator swing further, and by Eq. (5) larger theta swings modulate the gamma envelope more deeply.

The parameter *κ*_*a*_ has an additional significance. By Eq. (5), the gamma amplitude at the theta peak exceeds that at the trough by the factor 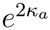. At C1 the median cycle has *κ*_*a*_ = 2.3, so the mean gamma envelope is about 100 times larger at the theta peak than at the trough.

## 4. Numerical methods

### 4.1. Simulation

We integrate Equation (3) using a semi-implicit scheme with a time step of 0.5 ms. By treating gamma friction and coupling together, this scheme preserves an exact property of the continuous equations: the total contraction rate of small gamma disturbances (Supplementary Section S2) [35]. Every record starts from the steady state of the uncoupled oscillators, runs 30 s with coupling to remove transients (burn-in), and is analysed over the following 10 s unless stated otherwise. Records with the same seed share initial state and noise across parameter values, so that comparisons between couplings are paired. All simulations and analyses were run in MATLAB (R2026a) with MATLAB’s default pseudorandom number generator (Mersenne twister), so that each seed reproduces the same noise. Table 2 lists all settings and seeds.

**Table 2:** Simulation settings.

| Quantity | Value | Use |
|---|---|---|
| Step, noise intensity | *h* = 0.5 ms, *σ* = 4.472 | all simulations |
| Frequencies | 4 Hz, 60 Hz | theta, gamma |
| Start, burn-in, record | uncoupled steady state, 30 s, 10 s | PAC analyses |
| Parameter scan | *g*<sub>*vv*</sub> = 0:0.1:10, *ζ* = 0.0002:0.0001:0.0055, seeds 100–199 | Sections 5.3 and 5.4 |
| Record-level subgrid | *g*<sub>*vv*</sub> = 0.5:0.5:10, 27 damping ratios, seeds 100–199 | Section 5.3 |
| Validation | seeds 4101–4120, 999 surrogates | Section 5.5 |
| Ten-term test | *ζ* = 0.0027, equal force strength, seeds 100–149 | Appendix B |
| Mixed friction terms | *ζ* = 0.0027, seeds 100–199 | Supplementary Section S4 |
| Other coupling terms | *g*<sub>*vv*</sub> > 0, *g*<sub>*vv*</sub> < 0, *g*<sub>*xv*</sub> > 0, *g*<sub>*xv*</sub> < 0, *K*<sub>*xx*</sub>, linear cross term; seeds 100–149 | Section 5.6 |

### 4.2. Measuring PAC

Theta phase and gamma amplitude are the instantaneous phase and envelope of the simulated coordinates, obtained with the standard analytic-signal method (Hilbert transform). Because the oscillator frequencies are fixed, no band-pass filter is needed for Eq. (3). The gamma amplitude is divided by one fitted exponential trend, which removes slow changes of scale over the record (detrending). For the modulation index, the theta cycle is divided into *N*_b_ = 18 phase bins, the mean gamma amplitude in each bin is normalized to a probability distribution, and MI is its distance from the uniform distribution (Supplementary Section S6) [8, 9]. The preferred phase is the theta phase at which this profile is centred, and zero is the theta peak. In the ten-term test, some terms add components far from the gamma frequency, so the gamma signal is first restricted to 40–80 Hz.

### 4.3. Simulation experiments

#### Fluctuation–dissipation tests

For theta, 200 records of 200 s give the effective temperature from the kinetic energy, from the potential energy and from the response to a brief push, which equipartition and the FDT predict to be equal (Supplementary Section S3). For gamma, 20 records of 200 s per coupling give the velocity variance, which is compared with Supplementary Eq. (S10); the response to a brief push, propagated along the same theta paths, gives a frequency-resolved temperature near 60 Hz, which would equal the kinetic temperature in equilibrium [36].

#### Parameter scan

The scan covers 101 ×54 combinations of coupling and damping ratio, with 100 records each. A combination is included when at least 95 of its records stay below a numerical ceiling of 10^12^ and when the coupling lies below 804.7 *ζ*^0.628^, a fit to the lower 95% confidence bound of the stability boundary measured in Part II [37]. Each included combination is summarized by its median MI.

#### Record-level test

To test Eq. (6) record by record, the scan is repeated on a coarser grid (*g*_*vv*_ = 0.5:0.5:10, 27 damping ratios), storing each record’s mean theta amplitude *ā*_*θ*_, from which its own *κ*_*a*_ = *g*_*vv*_*ā*_*θ*_*/*2 follows; 51,095 of 54,000 records are complete and included.

#### Validation

At C1 and at two neighbouring points, C2 = (8.1, 0.0028) and C3 = (8.175, 0.0027), we test whether the measured PAC is caused by the coupling. A surrogate test shifts the gamma envelope at random within each theta cycle, which destroys its phase relation to theta while preserving everything else; the observed MI must exceed that of 999 such surrogates [38], with *p* values corrected for testing three points [39]. An ablation test repeats each record with the coupling switched off but the same noise; MI must then drop. Records in which any variable exceeds 10^6^, a lower ceiling than the 10^12^ used in the scan and the ten-term test, are counted as incomplete and as failures, which makes the pass rates conservative. Pass rates count every record, with confidence intervals for a proportion (Wilson score intervals) [40].

#### Ten-term test

To compare the ten second-order terms fairly, each is added alone to the uncoupled gamma equation with a coefficient chosen so that its typical force equals that of the model’s coupling at C1 (*g*_*vv*_ = 8.2, *ζ* = 0.0027). Typical forces are root-mean-square values computed from the uncoupled steady state. For each term, 50 records (seeds 100–149) are simulated, and records in which any variable exceeds 10^12^ are excluded. We report the median MI and the preferred phase, together with how consistent that phase is across records (the length of the mean of the phases taken as unit vectors, 1 for identical phases and near 0 for random ones). Combinations of the two friction-type terms are tested with 100 records (seeds 100–199). The results are given in Appendix B.

#### Other coupling terms

To compare different coupling mechanisms, we modify the gamma equation of Eq. (3) with four kinds of coupling, six cases in all: *g*_*vv*_ = ±8.2; *g*_*xv*_ = ±8.2 *ω*_*θ*_; a stiffness-type term *K*_*xx*_*x*_*θ*_*x*_*γ*_ with the equal-force coefficient used in the ten-term test; and *g*_*vv*_ = 8.2 combined with a linear cross term −*g*_21_*v*_*θ*_ (*g*_21_ = 1000). The theta equation remains unchanged in all cases. For each case, we simulate 50 records (seeds 100–149) and measure PAC as in the ten-term test (Section 4.3), restricting the gamma signal to 40–80 Hz. We also estimate the instantaneous gamma frequency from the phase of the band-passed gamma signal and average it within theta-phase bins.

## 5. Results

The results below concern the velocity–velocity model of Eq. (3). The comparison of all ten second-order terms, which shows that only the friction-type terms produce PAC, is given in Appendix B.

### 5.1. PAC at weak and moderate coupling

Figure 2 illustrates the model along one theta path (seed 4120). At weak coupling (*g*_*vv*_ = 0.5) the gamma envelope shows no theta-locked structure (MI 0.001). At moderate coupling (*g*_*vv*_ = 8.2) gamma bursts at every theta peak and its logarithmic envelope rises and falls once per cycle, as Eq. (5) predicts (MI 0.20).

**Figure 2:**
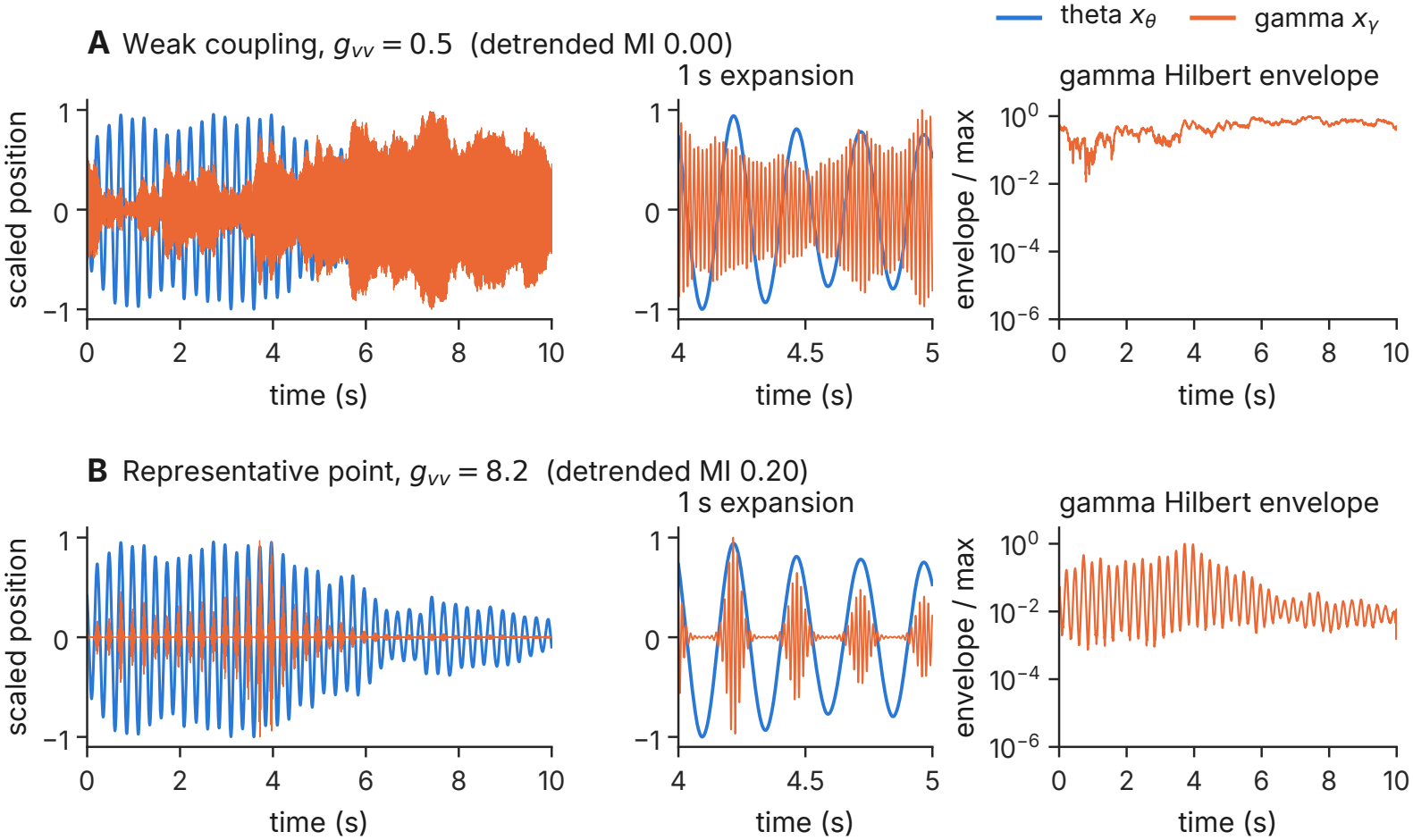
PAC at weak and moderate coupling. (*ζ* = 2.7 ×10^*−*3^, seed 4120). (A, B) *g*_*vv*_ = 0.5 and 8.2: scaled theta (blue) and gamma (orange) coordinates over a 10 s record after a 30 s burn-in, a 1 s expansion, and the gamma envelope on a logarithmic axis.

### 5.2. The fluctuation–dissipation relations hold

The fluctuation–dissipation relations on which the theory rests are confirmed directly (Fig. 3). For theta, three independent estimates of the effective temperature, from its kinetic energy, potential energy and response to a brief push, give 152.1, 152.1 and 152.2, each with a 95% interval that contains the predicted 147.4, and the response and the correlation function agree in shape to within 0.14%. For gamma, the measured variance follows the predicted growth with coupling to within 20% for *g*_*vv*_ ≤3 (Fig. 3; Supplementary Table S4). Its effective friction falls to about 0.4% of the baseline at *g*_*vv*_ = 4 (predicted 0.3%) but stays positive, as predicted for every finite coupling within the approximation of Supplementary Section S5. The coupling drives gamma out of equilibrium. Its temperature inferred from the response to a brief push near 60 Hz equals the kinetic temperature without coupling, but falls increasingly below it as the coupling grows, to about a ninth at *g*_*vv*_ = 4 (Supplementary Table S4). In equilibrium the two would be equal; the gap reflects the energy that the theta-modulated friction pumps into gamma.

**Figure 3:**
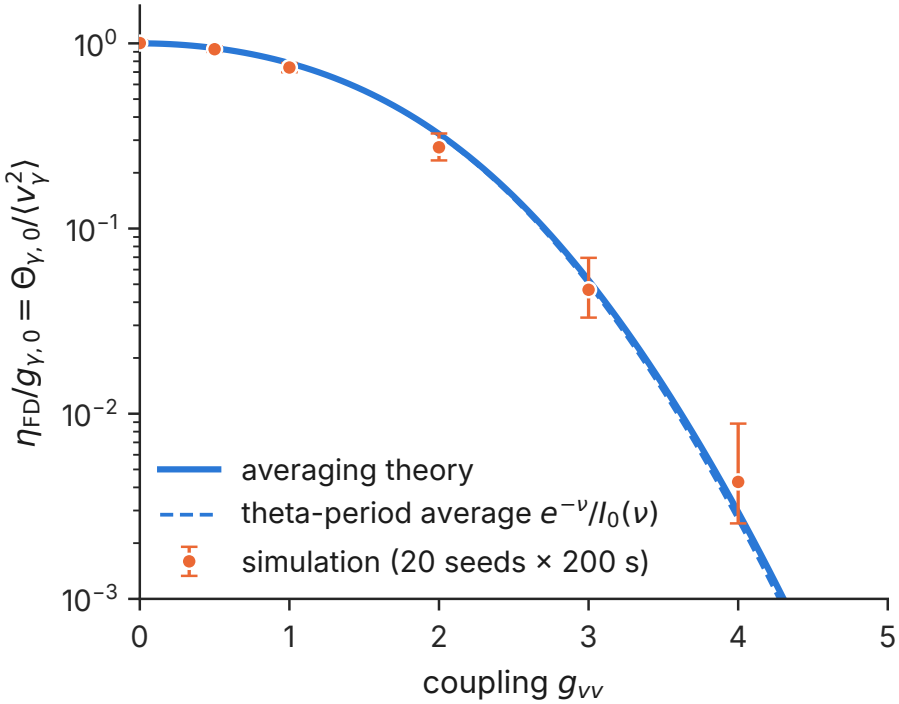
Fluctuation–dissipation relation of gamma. (*ζ* = 0.0027). Effective friction of gamma relative to its baseline, 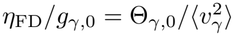, where Θ_*γ*,0_ = *σ*^2^*/*(2*g*_*γ*,0_) is the gamma temperature without coupling, against coupling: the prediction of Supplementary Eq. (S10) (solid), its theta-cycle average (dashed), and simulations (Supplementary Table S4). The friction decreases but stays positive.

### 5.3. One number predicts the PAC strength

The envelope law holds record by record: the slope of the logarithmic gamma amplitude against theta coordinate is within 2% of the predicted *g*_*vv*_*/*2 for couplings from 1 to 8.2 and 11% above it at *g*_*vv*_ = 0.5, where the modulation is weakest (seeds 4101–4120). Figure 4A plots the MI of 51,095 individual records against each record’s own *κ*_*a*_. They fall on the predicted curve Eq. (6), which explains 99.3% of their variance with no fitted parameter. Figure 4B plots the median MI of all 5,160 parameter combinations with *g*_*vv*_ *>* 0 against *κ* computed from the parameters alone; Eq. (A.7) explains 98.9% of the variance. The medians lie slightly below the prediction, by at most 0.03 and most at large *κ*, a deviation of the size expected from the first-order averaging of Section 3 and from averaging the envelope within each phase bin (Appendix A.4). At C1 the median of 100 records is 0.222, against a prediction of 0.233. The simplified model thus predicts PAC strength from one number.

**Figure 4:**
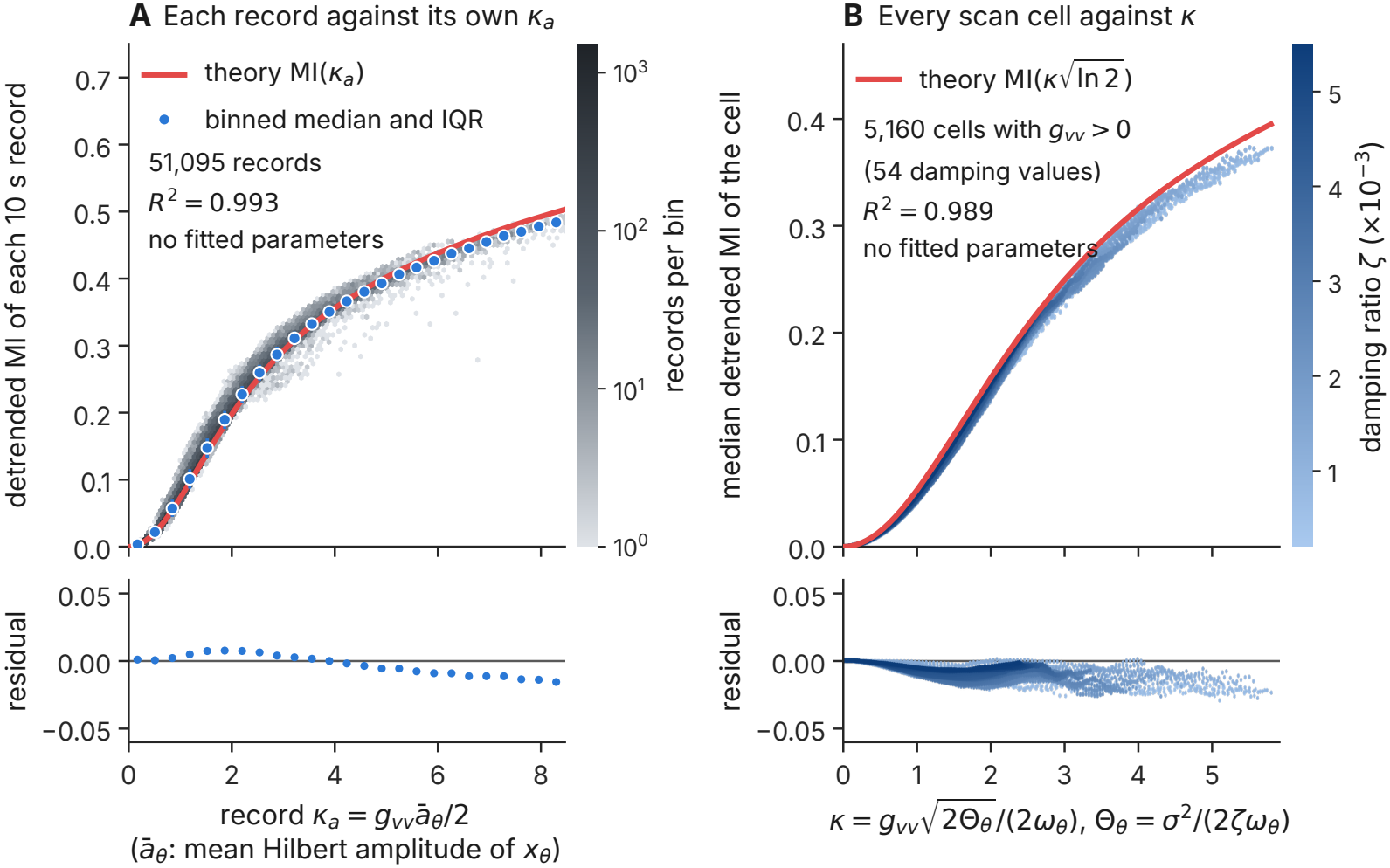
One number predicts the PAC strength. (A) MI of 51,095 10 s records against each record’s *κ*_*a*_ = *g*_*vv*_*ā*_*θ*_*/*2 (grey density), with binned medians and interquartile ranges (blue) and Eq. (6) (red). (B) Median MI of 5,160 parameter combinations against *κ*, coloured by damping ratio, with the prediction Eq. (A.7). Lower panels show residuals. No parameter is fitted.

### 5.4. PAC across the parameter plane

Across the included part of the parameter plane, MI rises steadily with coupling, and its contour lines follow lines of constant *κ* (Fig. 5). There is no interior optimum: the largest median MI, 0.372, lies on the edge of the included region. Of the 5,454 parameter combinations, 5,214 are included; all excluded ones lie at or beyond the bound used for inclusion (Section 4.3).

**Figure 5:**
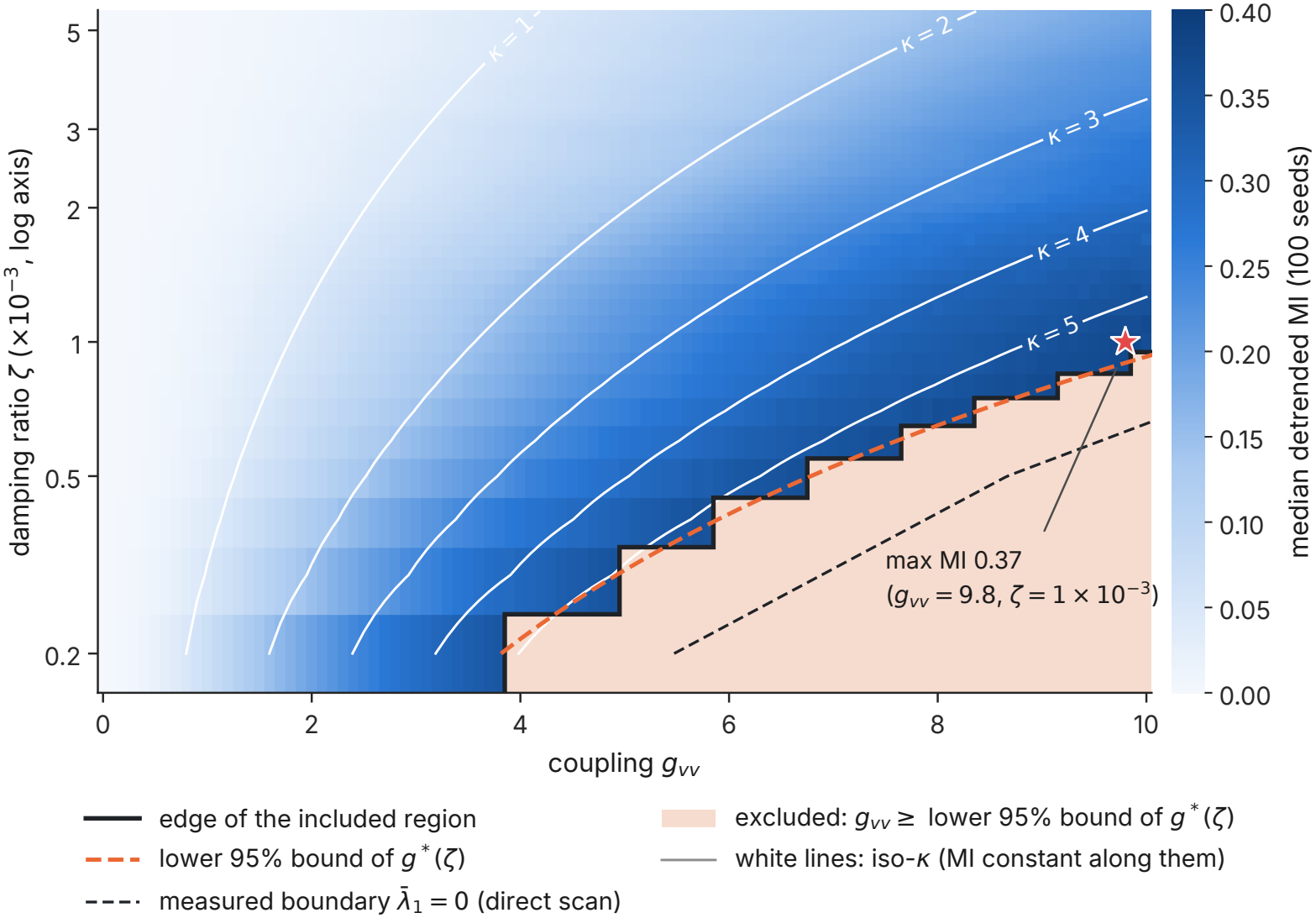
PAC across the parameter plane. Median MI of 100 records per combination of coupling and damping ratio, damping on a logarithmic axis. The black step line is the edge of the included region, the dashed orange curve the bound used for inclusion (the lower 95% bound of the boundary coupling *g*^*\**^(*ζ*) of Part II), and the dashed black curve the stability boundary measured in Part II, where the long-time growth rate of small gamma disturbances, 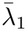, crosses zero. Shaded cells are excluded. White lines are lines of constant *κ*, along which MI is constant by Eq. (A.7). The star marks the largest median MI.

### 5.5. The coupling causes the measured PAC

At C1, C2 and C3, 17 of 20 records complete; the other three cross the numerical ceiling of 10^6^ during a large but bounded gamma burst (Part II [37]; Table 3). Every completed record exceeds all 999 surrogates, and switching the coupling off lowers MI to near zero, so PAC is caused by the coupling in 17 of 20 records at each point (95% interval 0.64–0.95). The preferred phase is −0.015 rad, the theta peak. The median MI of the 17 completed records at C1, 0.282 (bootstrap 95% interval 0.20–0.32), is consistent with the prediction 0.233.

**Table 3:** Validation at the representative point and two neighbours(seeds 4101–4120).

|  | *g*<sub>*vv*</sub> | *ζ* | Complete | Raw MI | Det. MI | Δ MI<sub>surr</sub> | Δ MI<sub>abl</sub> | PAC evidence |
|---|---|---|---|---|---|---|---|---|
| C1 | 8.200 | 0.0027 | 17/20 | 0.282 | 0.282 | 0.274 | 0.282 | 17/20 |
| C2 | 8.100 | 0.0028 | 17/20 | 0.276 | 0.276 | 0.250 | 0.276 | 17/20 |
| C3 | 8.175 | 0.0027 | 17/20 | 0.282 | 0.282 | 0.273 | 0.282 | 17/20 |

Medians over completed records. Det.: detrended. Δ MI<sub>surr</sub>: MI minus the surrogate median; Δ MI<sub>abl</sub>: MI minus that of the record without coupling. Prediction for C1: 0.233.

### 5.6. Simplifications of the model and other coupling terms

The model of Eq. (3) keeps the velocity–velocity term alone and rests on five simplifications, each of which can be tested:

a. *No memory, H* = 0, which holds if the hidden degrees of freedom forget faster than a gamma period. This is the strongest simplification (Section 6.2).
b. *No linear cross terms*: the two oscillators are far from resonance, so linear coupling leaves the gamma amplitude unchanged, although it adds a theta-frequency component to the gamma signal (tested below).
c. *No stiffness-type terms*: they would produce a theta-locked gamma frequency rather than a theta-locked gamma amplitude, which data can check (tested below).
d. *No influence of the gamma oscillator on the theta oscillator* : the theta oscillator is the pacing rhythm.
e. *Sinusoidal theta oscillations*: the theta oscillator has no nonlinearity of its own; non-sinusoidal waveforms are themselves a known source of spurious PAC [41].

Simplifications (b) and (c) concern terms that can be added back directly, and we do so here together with the other friction-type term. The velocity-velocity term is only one of the lowest-order interactions of Eq. (2). We did not analyse the others in as much detail, but sample trajectories show the range of behaviour (Fig. 6), and Table 4 summarizes what each kind of term does. At equal strength, *g*_*xv*_ = *g*_*vv*_*ω*_*θ*_, which gives the same *κ* by Supplementary Eq. (S8), the two friction-type terms produce the same PAC strength (median MI 0.20–0.21, 50 records each) but place the gamma bursts at different theta phases: *g*_*vv*_ *>* 0 at the peak (preferred phase −0.01 rad), *g*_*vv*_ *<* 0 at the trough (3.13 rad), *g*_*xv*_ *>* 0 on the falling phase (1.55 rad) and *g*_*xv*_ *<* 0 on the rising phase (− 1.59 rad) (Fig. 6A–E). A mixture of the two places them at any intermediate phase, *δ* = atan2(*g*_*xv*_*/ω*_*θ*_, *g*_*vv*_), with a PAC strength set by the combined size (Supplementary Section S4). Because recordings also show gamma activity at different theta phases [42, 43], the preferred phase alone does not single out the velocity–velocity term. The stiffness-type term *K*_*xx*_*x*_*θ*_*x*_*γ*_ instead makes the gamma frequency follow theta coordinate, swinging by 17 Hz over the theta cycle, while the gamma amplitude stays almost unmodulated (median MI 4 ×10^*−*4^; Fig. 6F): it produces phase– frequency coupling rather than PAC. A linear cross term, a force on the gamma oscillator proportional to theta velocity, adds a theta-frequency component to the gamma signal but leaves the PAC of the gamma band unchanged (median MI 0.215, preferred phase −0.01 rad). Because mixtures of terms can place the gamma bursts at the same theta phase, the terms have to be estimated jointly.

**Table 4:**
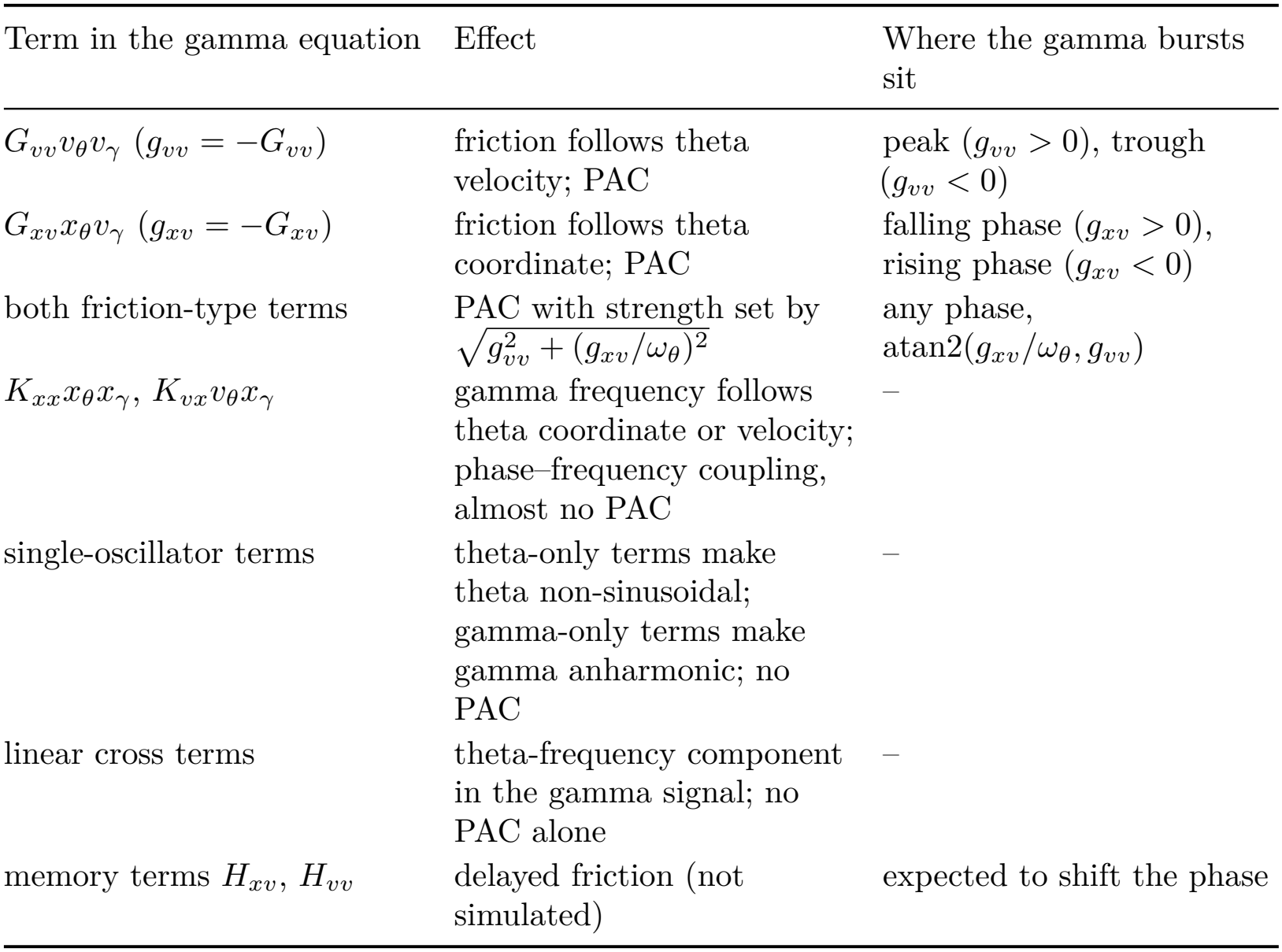
What each kind of lowest-order term does to the gamma oscillator.

| Term in the gamma equation | Effect | Where the gamma bursts sit |
|---|---|---|
| $G_{vv} v_\theta v_\gamma$ ($g_{vv} = -G_{vv}$) | friction follows theta velocity; PAC | peak ($g_{vv} > 0$), trough ($g_{vv} < 0$) |
| $G_{xv} x_\theta v_\gamma$ ($g_{xv} = -G_{xv}$) | friction follows theta coordinate; PAC | falling phase ($g_{xv} > 0$), rising phase ($g_{xv} < 0$) |
| both friction-type terms | PAC with strength set by $\sqrt{g_{vv}^2 + (g_{xv}/\omega_\theta)^2}$ | any phase, $\mathrm{atan2}(g_{xv}/\omega_\theta, g_{vv})$ |
| $K_{xx} x_\theta x_\gamma$, $K_{vx} v_\theta x_\gamma$ | gamma frequency follows theta coordinate or velocity; phase–frequency coupling, almost no PAC | – |
| single-oscillator terms | theta-only terms make theta non-sinusoidal; gamma-only terms make gamma anharmonic; no PAC | – |
| linear cross terms | theta-frequency component in the gamma signal; no PAC alone | – |
| memory terms $H_{xv}$, $H_{vv}$ | delayed friction (not simulated) | expected to shift the phase |

**Figure 6:**
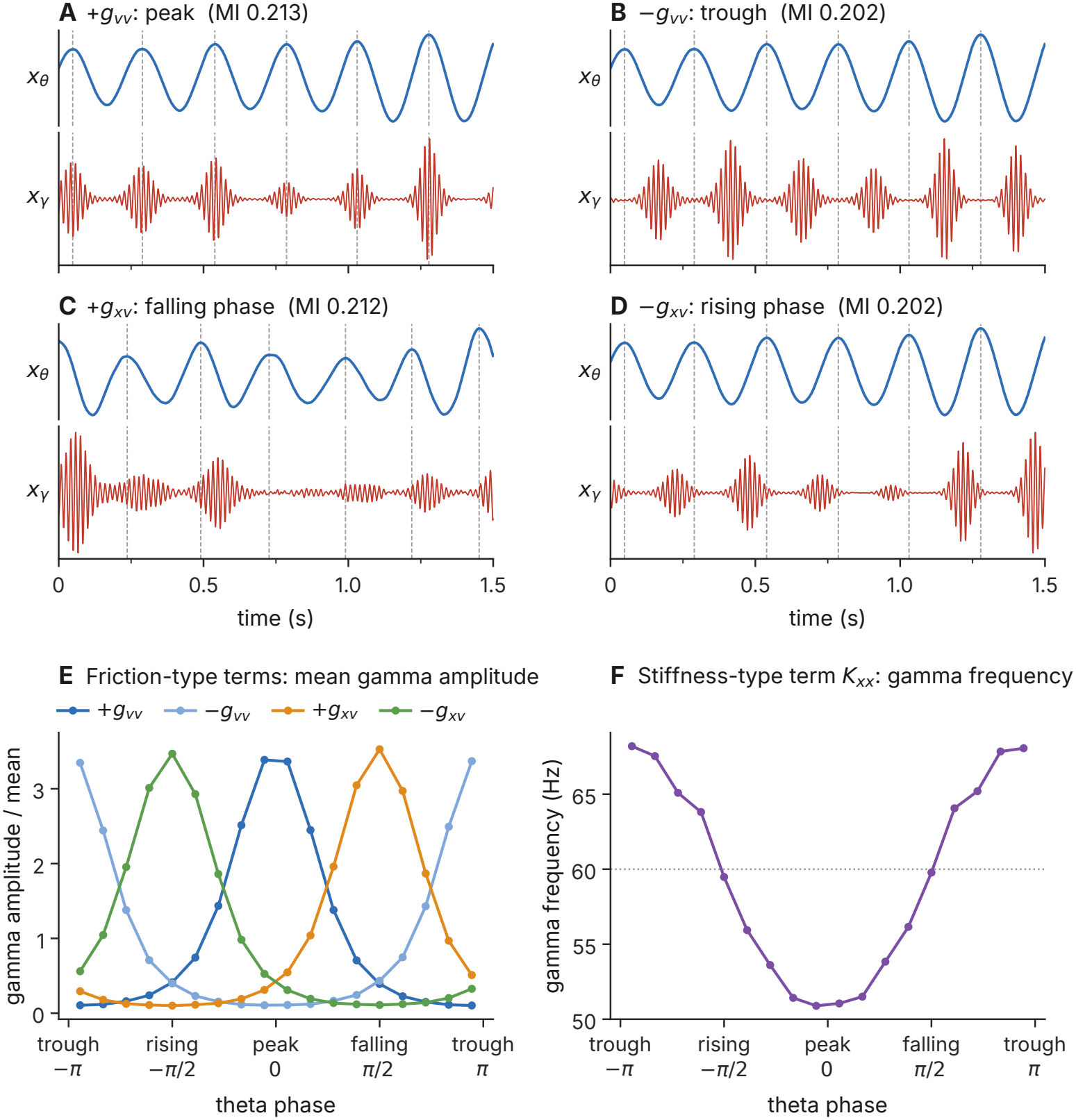
Other coupling terms. (*ζ* = 0.0027, strength of C1). (A–D) 1.5 s of theta (blue) and gamma (red, scaled to its maximum) in the record with the median MI among 50, for *g*_*vv*_ *>* 0, *g*_*vv*_ *<* 0, *g*_*xv*_ *>* 0 and *g*_*xv*_ *<* 0 (|*g*_*xv*_ |= |*g*_*vv*_| *ω*_*θ*_); dashed lines mark theta peaks. (E) Mean gamma amplitude (detrended, 40–80 Hz) against theta phase for the four friction-type cases. (F) Mean instantaneous gamma frequency against theta phase for the stiffness-type term *K*_*xx*_*x*_*θ*_*x*_*γ*_ at equal force strength.

## 6. Discussion

### 6.1. What this paper establishes

We asked whether a friction that follows theta velocity can produce theta– gamma PAC, and what then sets its strength. The question matters because PAC is widely measured and changes in epilepsy (Section 1), yet a value of MI alone does not say what produces it. To answer it, we chose the simplest system that has both rhythms: two damped oscillators driven by noise, with one coupling term. Like recordings, this system has theta and gamma rhythms whose amplitudes fluctuate at random. It is also simple enough that each result can be traced to one mechanism and derived in closed form.

A single friction term, with no fitted parameter, accounts for theta–gamma PAC. First, of the ten lowest-order ways in which theta can act on gamma, only the two friction-type terms produce PAC, and the term and its sign set the theta phase of the gamma bursts (Appendix B). Only a term that acts like a friction adds or removes gamma energy in every gamma cycle. The other terms change the gamma frequency or average out over a gamma cycle. Second, when the gamma friction follows theta velocity, the gamma amplitude follows theta coordinate as exp(*g*_*vv*_*x*_*θ*_*/*2) in every theta cycle. The reason is that the relative growth rate of the gamma amplitude is proportional to theta velocity, and adding up theta velocity over time gives the change in theta coordinate. Gamma therefore gains energy while theta rises and loses it while theta falls, so its bursts sit at the theta peak. Third, coupling, noise and damping set the PAC strength only through one number, 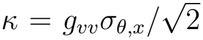. This number compares the coupling with the typical theta amplitude *σ*_*θ,x*_, so noise and damping matter only through how far theta typically swings. The closed form explains 99.3% of the variance of single records and 98.9% of the median PAC, so a measured MI becomes a measurement of *κ*. Fourth, the coupling drives gamma out of equilibrium. Theta keeps pumping energy into gamma, so the response of gamma to a brief push no longer matches its fluctuations, as it would in equilibrium. With the companion paper, in which the same *κ* sets the distance to a stability boundary [37], these results link the strength of PAC to the stability of the gamma rhythm.

Because Eq. (2) assumes no neuron model, it also gives existing models a common frame. Circuit, oscillator and neural-mass models in which theta modulates the excitability of the gamma population [17–21] act on gamma through a friction-type term, and regression and driven autoregressive models of PAC [44–46] estimate the friction-type terms, separating them from the stiffness-type terms that modulate only the gamma frequency.

### 6.2. Caveats and limitations

The model isolates *g*_*vv*_ rather than describing neural data, and it rests on the five simplifications of Section 5.6. The absence of memory is the strongest. Memory would add a delayed friction that could shift the preferred phase. Eq. (2) is an expansion, not a derivation from the microscopic dynamics, so its coefficients, such as *g*_*vv*_, are effective quantities. The theory also rests on well-separated time scales: gamma is fifteen times faster than theta, and both oscillators decay slowly compared with their periods (Table A.1 lists each approximation and its size). The main approximation, averaging over the fast gamma cycle, holds while gamma keeps oscillating within each theta cycle, that is, while the gamma friction stays well below critical damping. It also requires narrow spectral peaks. The damping ratios used here give peaks much narrower than those of EEG rhythms, and at EEG-like damping this approximation no longer holds. All results describe the steady state below the stability boundary of the companion paper [37]. In this state, the MI and the preferred phase do not depend on the initial coordinates and velocities, which are drawn at random for every record and then forgotten during the burn-in (Section 4.1). The other terms of Eq. (2) were tested alone, and *g*_*vv*_ was combined only with the other friction-type term and with a linear cross term. Combinations with stiffness-type and single-oscillator terms were not simulated. By the averaging argument of Supplementary Section S2, these terms should change the MI and the preferred phase only slightly, but this remains to be tested. The PAC estimator has known sensitivities to window length, transients and waveform shape [4, 41, 47–49], and a measured MI is also lowered by measurement noise, volume conduction and background activity. Finally, the preferred phase alone does not identify the mechanism, because different terms and their mixtures can place the gamma bursts at the same theta phase (Section 5.6).

### 6.3. Clinical relevance and future directions

Altered phase–amplitude coupling has been implicated in several neurological disorders, including enhanced beta–gamma coupling in Parkinson’s disease and reduced theta–gamma coupling in Alzheimer’s disease. This has motivated efforts to modulate PAC through external brain stimulation. Recent theoretical work using Stuart–Landau oscillators and coupled neural populations has shown how stimulation frequency, waveform, and phase can be optimized to enhance or suppress PAC [28]. Our approach provides a complementary perspective by relating PAC strength to effective friction, noise, and damping within a minimal stochastic dynamical model. Extending this framework to include memory effects and more realistic stimulation protocols could help identify the dynamical conditions under which stimulation enhances or suppresses PAC.

In the context of epilepsy, understanding the relationship between PAC and dynamical stability is particularly important.Theta–gamma coupling changes early in seizures and during epileptogenesis, and stimulation at theta frequencies can provoke seizures (Section 1). The model links these observations to quantities that recordings measure. Theta-band power gives the theta effective temperature up to the gain of the recording, and the theta spectral linewidth gives the theta friction. The gain cancels in the dimensionless *κ*, so that, given an observation model, a measured MI fixes *κ* through Eq. (A.7). The companion paper shows that the same *κ* also sets the distance of the gamma oscillator to a stability boundary [37], so a measured PAC could in principle indicate how close a recorded network is to instability. Whether this boundary corresponds to seizure onset is not tested here.

Three steps follow, each addressing a limitation above. First, because all coefficients of Eq. (2) enter linearly, the full equation, with memory, linear cross terms and all quadratic terms, can be fitted by linear regression on time-lagged coordinates and their products, which we will pursue in intracranial recordings. Such a fit would also show which terms act together in real data. The same recordings can test the simplifications of the model, by looking for a theta-locked gamma frequency (stiffness-type terms) and a gamma-dependent theta amplitude (an influence of gamma on theta). Second, relating a measured MI to *κ* requires an observation model, for which data-driven generalized Langevin methods offer a starting point [50]. Third, the theory should be extended to EEG-like damping and to memory terms, which the companion paper suggests are needed for a transition into seizure [37].

## Supporting information

Supplementary math

## Author Contributions

**Yuhan Luo:** Methodology, Software, Formal analysis, Investigation, Visualization, Writing – original draft. **Sonali Priyadarshini Nayak:** Methodology, Writing – review & editing. **Murielle Hsu:** Conceptualization, Writing – review & editing. **Gregory A. Worrell:** Conceptualization, Writing – review & editing. **John E. Straub:** Methodology, Writing – review & editing. **David Hsu:** Conceptualization, Software, Supervision, Project administration, Writing – original draft, Writing – review & editing.

## Data availability

The core simulation and analysis code (MATLAB) and the figures are available on GitHub at https://github.com/YuHanLuo-33/Velocity-dependent-friction-in-a-simple-two-oscillator-system-phase-ampli-tude-coupling-and-regions-.

## Declaration of generative AI and AI-assisted technologies in the manuscript preparation process

During the preparation of this work, the authors used Claude (Anthropic) for help in writing a first draft of the Introduction, and OpenEvidence to help with literature searches. AI was not used for the main body of the manuscript, nor in the writing of the computer codes, nor in analysis.

## Appendix A. Derivation of the envelope law and the PAC closed form

This appendix derives Eqs. (A.7), (5) and (6) in four steps, each building on the one before. The theta statistics of Eq. (4) and the distribution of theta amplitudes (a Rayleigh distribution) are derived in Supplementary Section S3.

1. The gamma amplitude obeys an exact equation, in which the friction acts only on the kinetic share of the gamma energy (Appendix A.1).
2. Averaging over the fast gamma phase replaces the kinetic share by one half, and the theta velocity in the friction integrates to a displacement. This gives the envelope law (Appendix A.2).
3. The gamma noise sets the overall gamma level, while the present theta coordinate sets the shape of the envelope within a theta cycle (Appendix A.3).
4. This shape is a single smooth peak on the theta cycle, whose modulation index has a closed form. Its median over records follows from the median theta amplitude (Appendix A.4).

The rates of the model are well separated, *ω*_*γ*_≫ *ω*_*θ*_ ≫*g*_*γ*,0_ ≫*g*_*θ*,0_ (377, 25, 1.0 and 0.07 s^*−*1^ at *ζ* = 0.0027). Gamma oscillates fifteen times per theta cycle, it forgets its energy over about four theta cycles, and the theta amplitude changes more slowly still. Approximate steps are marked by ≈or *O* (*ε*). The small parameter *ε* = max |*g*_*γ*,eff_| */*(2*ω*_*γ*_) compares the largest gamma friction in a theta cycle, *g*_*γ*,eff_ = *g*_*γ*,0_− *g*_*vv*_*v*_*θ*_, with critical damping, the friction 2*ω*_*γ*_ at which gamma stops oscillating. Table A.1 lists every approximation, where it enters and its size at C1. Supplementary Table S2 lists the symbols.

**Table A.1:**
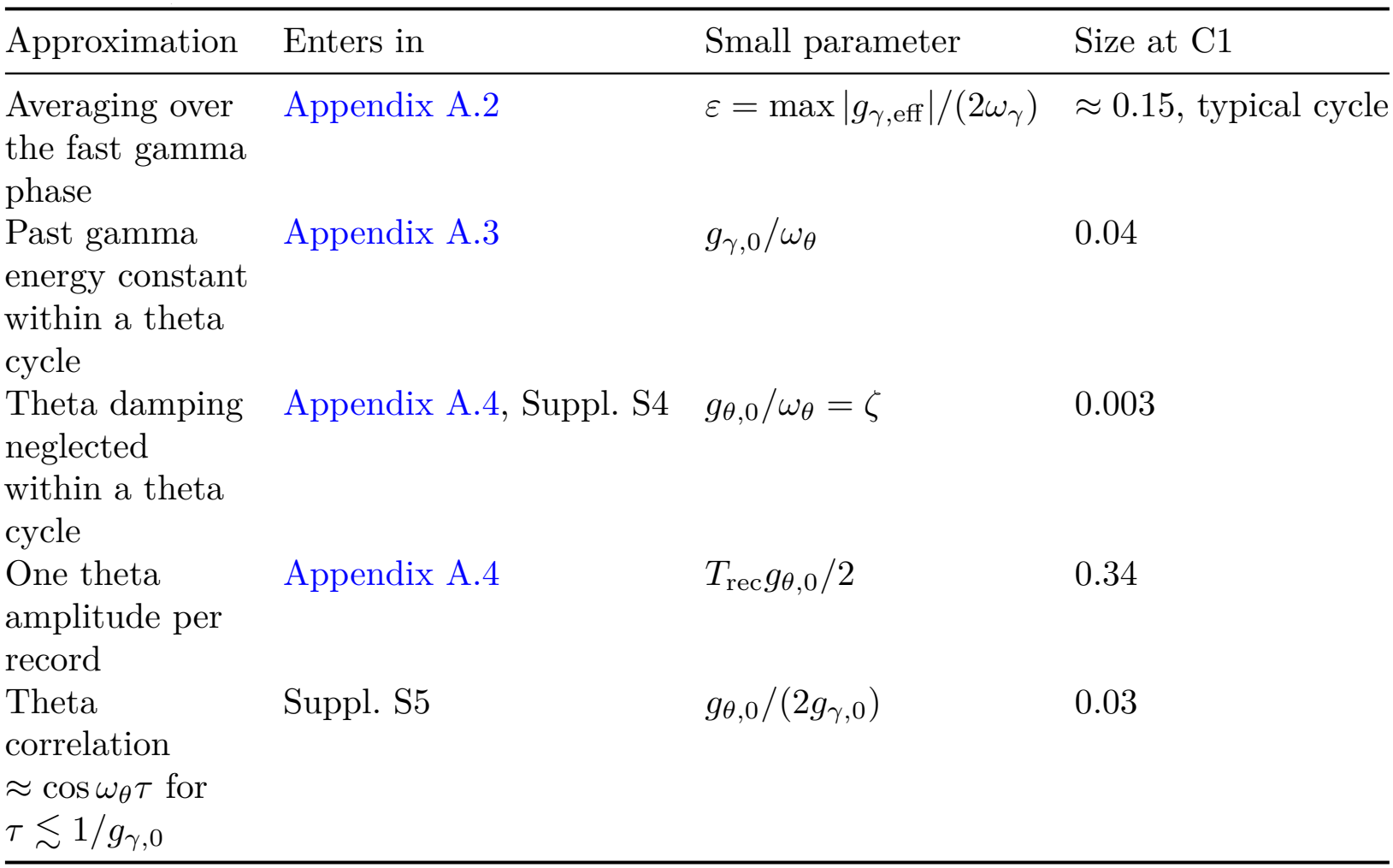
Approximations used in the derivations, with their size at C1 (*g*_*vv*_ = 8.2, *ζ* = 0.0027). *T*_rec_ = 10 s is the record length and *τ* a time lag.

| Approximation | Enters in | Small parameter | Size at C1 |
|---|---|---|---|
| Averaging over the fast gamma phase | Appendix A.2 | ε = max \|g<sub>γ,eff</sub>\|/(2ω<sub>γ</sub>) | ≈ 0.15, typical cycle |
| Past gamma energy constant within a theta cycle | Appendix A.3 | g<sub>γ,0</sub>/ω<sub>θ</sub> | 0.04 |
| Theta damping neglected within a theta cycle | Appendix A.4, Suppl. S4 | g<sub>θ,0</sub>/ω<sub>θ</sub> = ζ | 0.003 |
| One theta amplitude per record | Appendix A.4 | T<sub>rec</sub>g<sub>θ,0</sub>/2 | 0.34 |
| Theta correlation ≈ cos ω<sub>θ</sub>τ for τ ≲ 1/g<sub>γ,0</sub> | Suppl. S5 | g<sub>θ,0</sub>/(2g<sub>γ,0</sub>) | 0.03 |

### A.1. Gamma: exact amplitude and phase equations

Along a given theta path, and without noise (noise is added in Appendix A.3), gamma obeys 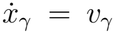 and 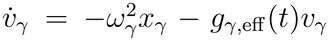, with *g*_*γ*,eff_(*t*) = *g*_*γ*,0_ − *g*_*vv*_*v*_*θ*_(*t*). We write gamma in amplitude–phase variables,

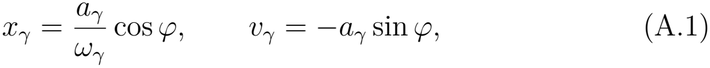

where *r* ≥ 0 is the gamma amplitude in velocity units and *φ* is the fast gamma phase, which advances at about *ω*_*γ*_. Then 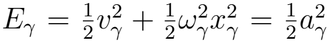 is the gamma energy, and sin^2^ 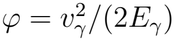 is its kinetic share, the fraction of the energy carried by the velocity. Because *a*_*γ*_ changes slowly compared with *φ*, the envelope of *x*_*γ*_ used in Section 4.2 (its Hilbert envelope) is *A*_*γ*_ ≈*r/ω*_*γ*_. Differentiating Eq. (A.1) in time, substituting the equations of motion and solving for 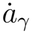 and 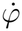 gives

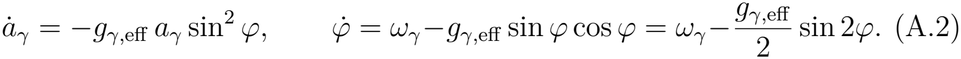

These equations are exact. Multiplied by *a*_*γ*_, the first is the energy balance 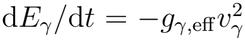. The friction thus acts only on the kinetic share of the gamma energy, whereas the spring only moves energy between coordinate and velocity and does no net work. The second equation shows that the friction also speeds up or slows down the rotation of the phase. The envelope law needs only one approximation, made in Appendix A.2, where the kinetic share is replaced by its average.

For a constant friction *g*, the phase stops rotating where 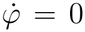, that is, where sin 2*φ* = 2*ω*_*γ*_*/g*. Because |sin 2*φ*| ≤1, such phases exist only if |*g*| ≥2*ω*_*γ*_. Below this friction, called critical damping, the phase keeps rotating and gamma oscillates. Above it, the phase locks and gamma grows or decays without oscillating. The roots of the characteristic equation 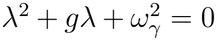 of 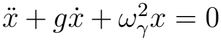 show the same: they are complex below critical damping and real above it. Part II [37] uses this to derive the stability boundary.

### Appendix A.2. Averaging: the envelope law

This step proves Eq. (5). By Eq. (A.2) and sin^2^ 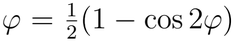, the logarithm of the gamma amplitude changes as

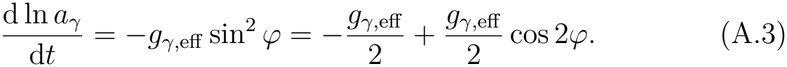

The second term oscillates at about 2*ω*_*γ*_. We show that it contributes nothing for a constant friction and only a small, bounded correction for a friction that follows theta.

i. *Constant friction*. Let the friction be a constant *g* with |*g* |*<* 2*ω*_*γ*_, so that the phase keeps rotating. The phase spends the time 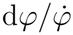 near each value *φ*, so the time average of cos 2*φ* over one turn is proportional to ∮ cos 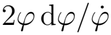. With *u* = sin 2*φ*, cos 2*φ* d*φ* = d*u/*2 and 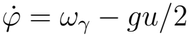, this integral becomes ∮ d*u/*(2*ω*_*γ*_− *gu*). It vanishes, because *u* returns to its starting value after each turn and the integrand is a smooth function of *u*. Hence the kinetic share averages to exactly 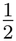, and ln *a*_*γ*_ changes at the rate −*g/*2, the real part of the characteristic roots.
ii. *Friction that follows theta*. Now let *g*_*γ*,eff_ change on the theta time scale. Writing cos 2*φ* d*t* = d(sin 2*φ*)*/*(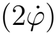) and integrating by parts gives

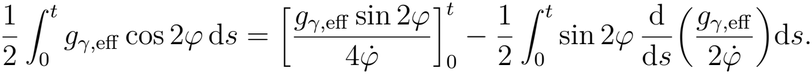

By Eq. (A.2), 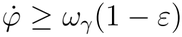, so each end of the boundary term is at most *ε/*[2(1 *ε*)] in size, and the term is *O*(*ε*). In the remaining integral, the derivative of 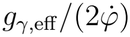 is of order *εω*_*θ*_ for its slow part and of order *ε*^2^*ω*_*γ*_ for the part that oscillates with the gamma phase. Multiplied by sin 2*φ* and integrated, both give bounded terms that are smaller than the boundary term by a factor *ω*_*θ*_*/ω*_*γ*_ or *ε*. Neither term grows with *t*, so the oscillating part of Eq. (A.3) adds to ln *a*_*γ*_ only a bounded correction of order *ε*, which does not accumulate.

#### The envelope law

Integrating Eq. (A.3) therefore gives ln 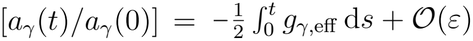 integral of a velocity is a displacement, 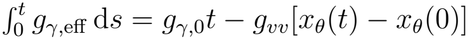. Theta enters only through its velocity, and the time so that

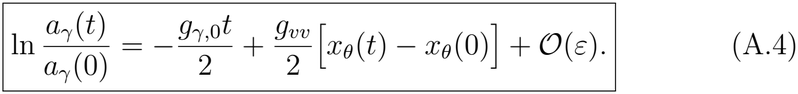

Equation (A.4) holds along every theta path, without averaging over theta. The gamma amplitude is therefore exp(*g*_*vv*_*x*_*θ*_*/*2) times a factor that decays at the baseline rate *g*_*γ*,0_*/*2, and it is largest at the theta coordinate peak. For the coordinate-dependent friction term *x*_*θ*_*v*_*γ*_, the same integral contains *∫x*_*θ*_ d*s* instead. Within a theta cycle this is proportional to minus the theta velocity, which moves the envelope peak a quarter cycle later (Supplementary Section S4). The derivation requires *ε <* 1, that is, that gamma keeps oscillating. Part II [37] treats the rare theta cycles in which this fails.

### Appendix A.3. Noise sets the level, theta coordinate sets the shape

Equation (A.4) says how a given gamma amplitude evolves, but without noise gamma would decay to zero. The noise sets the level at which the energy it injects balances the energy that friction removes. With noise, the chain rule for noise-driven variables (Itô’s formula) gives the gamma energy equation

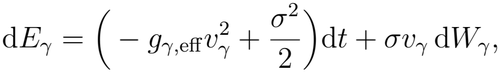

where *W*_*γ*_ is the accumulated gamma noise, d*W*_*γ*_ = *ξ*_*γ*_ d*t* (a Wiener process). Compared with the ordinary chain rule, Itô’s formula adds the term *σ*^2^*/*2, the mean power that the noise injects. We now average over the gamma noise along a fixed theta path and denote this average by ⟨·⟩_*θ*_. The noise term *σv*_*γ*_ d*W*_*γ*_ then has zero mean. Averaging over the fast gamma phase, as in Appendix A.2, replaces *v*^2^ = 2*E*_*γ*_ sin^2^ *φ* by *E*_*γ*_. This gives a linear equation for ⟨*E*_*γ*_ ⟩_*θ*_, which we solve by the standard integrating-factor method, starting from the distant past so that the initial state is forgotten:

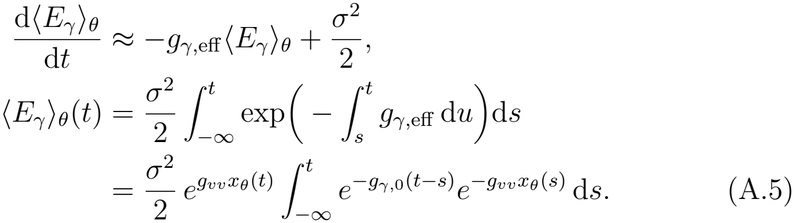

The last line uses 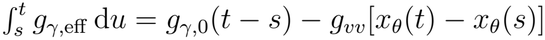, which is again exact because the time integral of theta velocity is a displacement. The first factor, 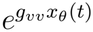, depends only on the present theta coordinate. The remaining integral is a weighted average of 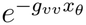 over the past 1*/g*_*γ*,0_ ≈ 1 s, about four theta cycles. Because *g*_*γ*,0_ ≪ *ω*_*θ*_, it changes little within one theta cycle (Table A.1). Within a cycle, therefore, 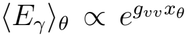. Since 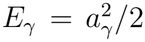 and *A*_*γ*_ ≈ *r/ω*_*γ*_, the typical gamma envelope, 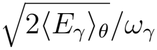, is proportional to 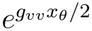. The noise thus sets the overall level, and the theta coordinate sets the shape.

### Appendix A.4. From the envelope shape to the modulation index

This step proves Eqs. (A.7) and (6).

*The envelope over one theta cycle*. Within one theta cycle the theta amplitude *a*_*θ*_ hardly changes (Table A.1), so *x*_*θ*_ ≈ *a*_*θ*_ cos *ϕ*_*θ*_. Here *ϕ*_*θ*_ is the theta phase of Section 4.2 (its Hilbert phase), zero at the coordinate peak. Inserted into the result of Appendix A.3, this gives *A*_*γ*_ ∝ exp 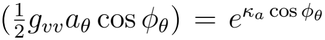, with 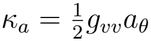. The modulation index treats the envelope, normalized over theta phase, as a probability density *p*(*ϕ*_*θ*_) and compares it with a uniform density. Normalized, the envelope is a single smooth peak on the theta cycle whose sharpness is set by *κ*_*a*_ (a von Mises density):

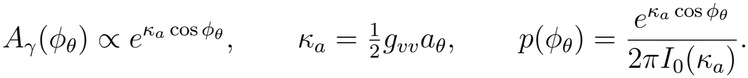

The normalization uses 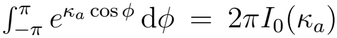. Here *I*_0_ and *I*_1_, used below, are standard functions available in numerical libraries (modified Bessel functions of the first kind).

#### Modulation index

The modulation index measures how far *p* departs from the uniform density 1*/*(2*π*) (the Kullback–Leibler divergence), divided by ln *N*_b_, where *N*_b_ = 18 is the number of phase bins (Supplementary Section S6). For the density above, ln(2*πp*) = *κ*_*a*_ cos *ϕ*_*θ*_ ln *I*_0_ (*κ*_*a*_). The mean of cos *ϕ*_*θ*_ is 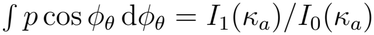, because d*I*_0_*/*d*κ*_*a*_ = *I*_1_. Hence

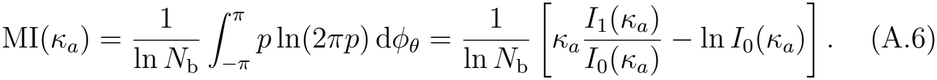

For small 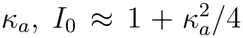 and *I*_1_ ≈ *κ*_*a*_*/*2, so MI 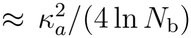 grows quadratically. For large *κ*_*a*_, *I*_1_/ *I*_0_≈ 1−1/(2*κ*_*a*_) and ln 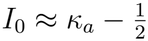 ln(2 *πκ*_*a*_), so MI ≈ ln(2*πκ*_*a*_*/e*)*/*(2 ln *N*_b_) grows only logarithmically. If the envelope is evaluated at the bin centres, the binned index with *N*_b_ = 18 agrees with Eq. (A.6) to better than 10^*−*6^ for *κ*_*a*_ ≤8. The estimator averages the envelope over each bin instead, which lowers it slightly: by 0.001 at *κ*_*a*_ = 1, 0.003 at *κ*_*a*_ = 2.3 (C1) and 0.013 at *κ*_*a*_ = 8 [9, 51].

#### Median over records

The theta amplitude changes over 2*/g*_*θ*,0_≈ 30 s, so a 10 s record contains essentially one theta amplitude, drawn from the Rayleigh distribution of Supplementary Section S3. Because MI increases monotonically with *a*_*θ*_, the median of MI over records is its value at the median theta amplitude, 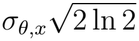:

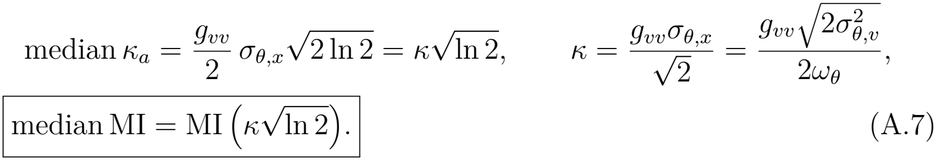

Section 3.3 discusses its meaning.

## Appendix B. The ten second-order terms at equal force strength

This appendix tests the classification of Section 2.2 directly. Each of the ten second-order terms is added alone to the uncoupled gamma equation, with a coefficient chosen so that its typical force equals that of the coupling at C1 (Section 4.3). Figure B.1 shows a representative record for each term, and Supplementary Table S1 lists all values. Only the two friction-type terms produce clear PAC, with median MI 0.213 for *v*_*θ*_*v*_*γ*_ and 0.212 for *x*_*θ*_*v*_*γ*_, close to the predicted 0.233. The two stiffness-type terms give MI 4× 10^*−*4^, about five hundred times weaker, because they change the gamma frequency but not the gamma energy (Supplementary Section S2). The six single-oscillator terms leave gamma unmodulated (MI 6 ×10^*−*5^ to 1 ×10^*−*4^). The two friction-type terms differ in where gamma is locked: at the theta peak for *v*_*θ*_*v*_*γ*_ (preferred phase −0.014 rad) and a quarter cycle later for *x*_*θ*_*v*_*γ*_ (1.553 rad). Their mixtures behave as predicted (Supplementary Table S3).

**Figure B.1:**
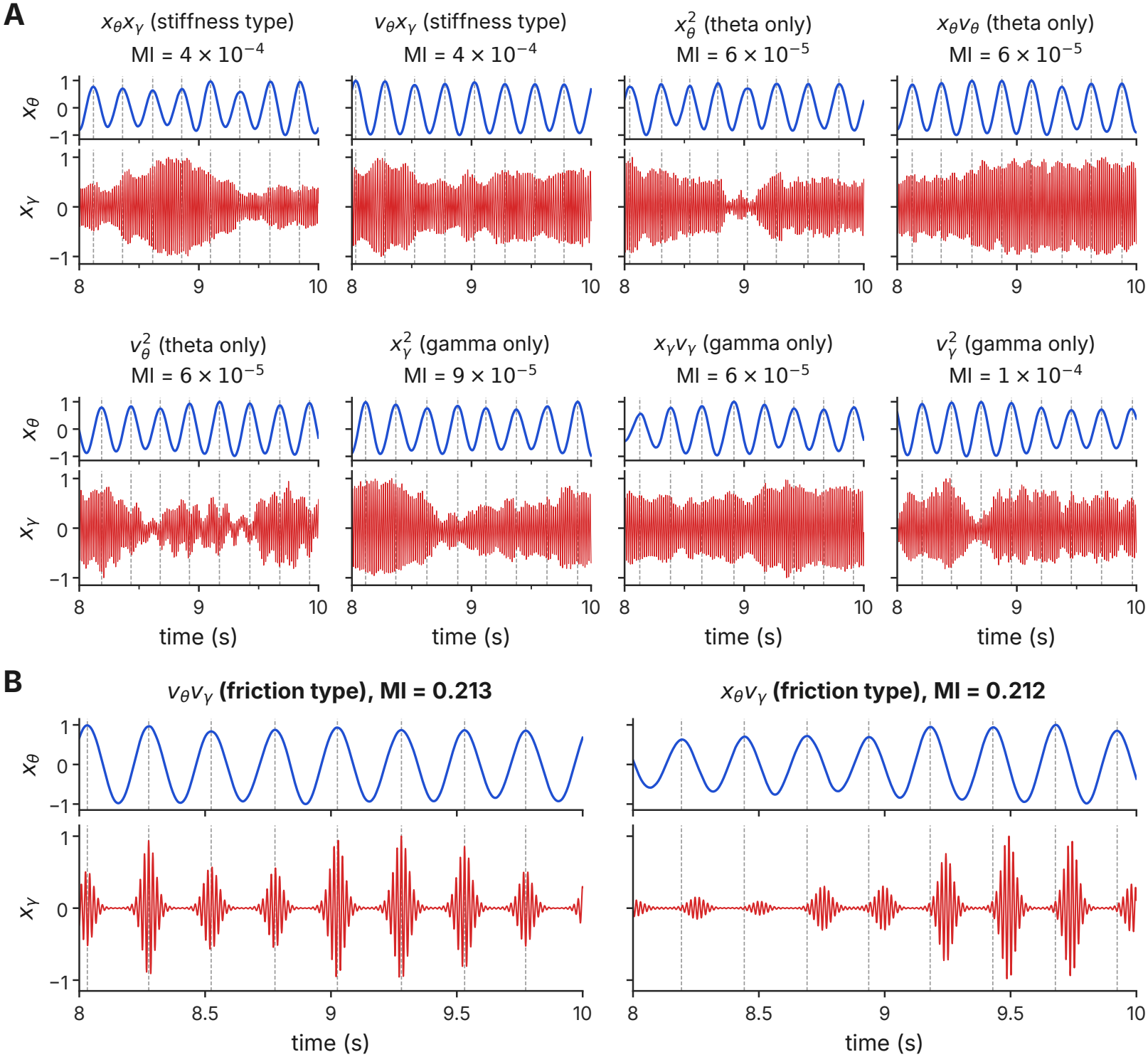
The ten second-order theta–gamma terms at equal force strength. (*ζ* = 0.0027). Each cell shows 2 s of the record with the median MI among 50: the theta coordinate (blue) above and the gamma coordinate (red) below, each scaled to its maximum. Grey dashed lines mark theta peaks. Titles give the term, its kind and the median MI. (A) The eight terms that are not of friction type: stiffness-type terms (*x*_*θ*_*x*_*γ*_, *v*_*θ*_*x*_*γ*_) and single-oscillator terms. (B) The two friction-type terms. Gamma bursts at the theta peaks for *v*_*θ*_*v*_*γ*_ and a quarter cycle later for *x*_*θ*_*v*_*γ*_.

## References

[1] O. Jensen, L. L. Colgin, Cross-frequency coupling between neuronal oscillations, Trends in Cognitive Sciences 11 (7) (2007) 267–269. doi: 10.1016/j.tics.2007.05.003.

[2] R. T. Canolty, R. T. Knight, The functional role of cross-frequency coupling, Trends in Cognitive Sciences 14 (11) (2010) 506–515.

[3] A. Hyafil, A.-L. Giraud, L. Fontolan, B. Gutkin, Neural cross-frequency coupling: Connecting architectures, mechanisms, and functions, Trends in Neurosciences 38 (11) (2015) 725–740. doi:10.1016/j.tins.2015.09.001.

[4] J. Aru, J. Aru, V. Priesemann, M. Wibral, L. Lana, G. Pipa, W. Singer, R. Vicente, Untangling cross-frequency coupling in neuroscience, Current Opinion in Neurobiology 31 (2015) 51–61. doi:10.1016/j.conb.2014.08.002.

[5] J. E. Lisman, M. A. P. Idiart, Storage of 7 plus or minus 2 short-term memories in oscillatory subcycles, Science 267 (5203) (1995) 1512–1515. doi:10.1126/science.7878473.

[6] J. E. Lisman, O. Jensen, The theta-gamma neural code, Neuron 77 (6) (2013) 1002–1016. doi:10.1016/j.neuron.2013.03.007.

[7] R. T. Canolty, E. Edwards, S. S. Dalal, M. Soltani, S. S. Nagarajan, H. E. Kirsch, M. S. Berger, N. M. Barbaro, R. T. Knight, High gamma power is phase-locked to theta oscillations in human neocortex, Science 313 (5793) (2006) 1626–1628. doi:10.1126/science.1128115.

[8] A. B. L. Tort, M. A. Kramer, C. Thorn, D. J. Gibson, Y. Kubota, A. M. Graybiel, N. J. Kopell, Dynamic cross-frequency couplings of local field potential oscillations in rat striatum and hippocampus during performance of a T-maze task, Proceedings of the National Academy of Sciences 105 (51) (2008) 20517–20522. doi:10.1073/pnas.0810524105.

[9] A. B. L. Tort, R. Komorowski, H. Eichenbaum, N. Kopell, Measuring phase-amplitude coupling between neuronal oscillations of different frequencies, Journal of Neurophysiology 104 (2) (2010) 1195–1210. doi:10.1152/jn.00106.2010.

[10] N. Axmacher, M. M. Henseler, O. Jensen, I. Weinreich, C. E. Elger, J. Fell, Cross-frequency coupling supports multi-item working memory in the human hippocampus, Proceedings of the National Academy of Sciences of the USA 107 (7) (2010) 3228–3233.

[11] P. Sauseng, W. Klimesch, K. F. Heise, W. R. Gruber, E. Holz, A. A. Karim, M. Glennon, C. Gerloff, N. Birbaumer, F. C. Hummel, Brain oscillatory substrates of visual short-term memory capacity, Current Biology 19 (21) (2009) 1846–1852.

[12] X. Liu, F. Han, R. Fu, Q. Wang, G. Luan, Epileptogenic zone location of temporal lobe epilepsy by cross-frequency coupling analysis, Frontiers in Neurology 12 (2021) 764821. doi:10.3389/fneur.2021.764821.

[13] S. A. Weiss, G. P. Banks, G. M. McKhann, R. R. Goodman, R. G. Emerson, A. J. Trevelyan, C. A. Schevon, Ictal high frequency oscillations distinguish two types of seizure territories in humans, Brain 136 (12) (2013) 3796–3808.

[14] A. Darves-Bornoz, E. J. Barbeau, M. Denuelle, A. Calvel, A. De Barros, Z. Darrasse, K. Guines, J.-A. Lotterie, L. Valton, J. Curot, Thetarange SEEG stimulation for temporal lobe mapping: an alternative to conventional 1-Hz and 50-Hz protocols, medRxiv preprint (2026). doi:10.64898/2026.03.31.26349175.

[15] N. A. Jansen, C. Perez, M. Schenke, A. W. van Beurden, A. Dehghani, R. A. Voskuyl, R. D. Thijs, G. Ullah, A. M. J. M. van den Maagdenberg, E. A. Tolner, Impaired θ-γ coupling indicates inhibitory dysfunction and seizure risk in a Dravet syndrome mouse model, Journal of Neuroscience 41 (3) (2021) 524–537. doi:10.1523/JNEUROSCI.2132-20.2020.

[16] D. Hsu, M. Hsu, H. L. Grabenstatter, G. A. Worrell, T. P. Sutula, Timefrequency analysis using damped-oscillator pseudo-wavelets: Application to electrophysiological recordings, Journal of Neuroscience Methods 194 (1) (2010) 179–192. doi:10.1016/j.jneumeth.2010.09.019.

[17] A. C. E. Onslow, M. W. Jones, R. Bogacz, A canonical circuit for generating phase-amplitude coupling, PLoS ONE 9 (8) (2014) e102591. doi:10.1371/journal.pone.0102591.

[18] Y. Qin, T. Menara, D. S. Bassett, F. Pasqualetti, Phase-amplitude coupling in neuronal oscillator networks, Physical Review Research 3 (2021) 023218. doi:10.1103/PhysRevResearch.3.023218.

[19] R. C. Sotero, Topology, cross-frequency, and same-frequency band interactions shape the generation of phase-amplitude coupling in a neural mass model of a cortical column, PLoS Computational Biology 12 (11) (2016) e1005180. doi:10.1371/journal.pcbi.1005180.

[20] M. Chehelcheraghi, C. van Leeuwen, E. Steur, C. Nakatani, A neural mass model of cross frequency coupling, PLOS ONE 12 (4) (2017) e0173776. doi:10.1371/journal.pone.0173776.

[21] M. Segneri, H. Bi, S. Olmi, A. Torcini, Theta-nested gamma oscillations in next generation neural mass models, Frontiers in Computational Neuroscience 14 (2020) 47. doi:10.3389/fncom.2020.00047.

[22] E. Montbrió, D. Pazó, A. Roxin, Macroscopic description for networks of spiking neurons, Physical Review X 5 (2015) 021028. doi:10.1103/PhysRevX.5.021028.

[23] V. K. Jirsa, W. C. Stacey, P. P. Quilichini, A. I. Ivanov, C. Bernard, On the nature of seizure dynamics, Brain 137 (8) (2014) 2210–2230.

[24] V. K. Jirsa, T. Proix, D. Perdikis, M. M. Woodman, H. Wang, J. Gonzalez-Martinez, C. Bernard, C. Bénar, M. Guye, P. Chauvel, F. Bartolomei, The virtual epileptic patient: individualized whole-brain models of epilepsy spread, NeuroImage 145 (2017) 377–388.

[25] M. Hashemi, A. N. Vattikonda, V. Sip, M. Guye, F. Bartolomei, M. M. Woodman, V. K. Jirsa, The Bayesian virtual epileptic patient: a probabilistic framework designed to infer the spatial map of epileptogenicity in a personalized large-scale brain model of epilepsy spread, NeuroImage 217 (2020) 116839.

[26] G. Buzsáki, C. A. Anastassiou, C. Koch, The origin of extracellular fields and currents—EEG, ECoG, LFP and spikes, Nature Reviews Neuroscience 13 (6) (2012) 407–420.

[27] P. L. Nunez, R. Srinivasan, Electric Fields of the Brain: The Neurophysics of EEG, 2nd Edition, Oxford University Press, New York, 2006.

[28] B. Duchet, R. Bogacz, How to design optimal brain stimulation to modulate phase-amplitude coupling?, Journal of Neural Engineering 21 (4) (2024) 046006. doi:10.1088/1741-2552/ad5b1a.

[29] S. P. Nayak, S. Roy, Immune phase transition under steroid treatment, Phys. Rev. E 103 (2021) 062401. doi:10.1103/PhysRevE.103.062401.

[30] S. P. Nayak, B. Bagchi, S. Roy, Effects of immunosuppressants on T-cell dynamics: Understanding from a generic coarse-grained immune network model, Journal of Biosciences 47 (4) (2022) 70. doi:10.1007/s12038-022-00312-4.

[31] S. Sahoo, S. P. Nayak, K. Hari, P. Purkait, S. Mandal, A. Kishore, H. Levine, M. K. Jolly, Immunosuppressive traits of the hybrid epithelial/mesenchymal phenotype, Frontiers in Immunology 12 (2021) 797261. doi:10.3389/fimmu.2021.797261.

[32] R. Kubo, The fluctuation-dissipation theorem, Reports on Progress in Physics 29 (1) (1966) 255–284. doi:10.1088/0034-4885/29/1/306.

[33] D. Hsu, M. Hsu, Zwanzig–Mori projection operators and EEG dynamics: Deriving a simple equation of motion, PMC Biophysics 2 (2009) 6. doi:10.1186/1757-5036-2-6.

[34] U. M. B. Marconi, A. Puglisi, L. Rondoni, A. Vulpiani, Fluctuation– dissipation: Response theory in statistical physics, Physics Reports 461 (2008) 111–195.

[35] P. E. Kloeden, E. Platen, Numerical Solution of Stochastic Differential Equations, Springer, Berlin, 1992.

[36] L. F. Cugliandolo, J. Kurchan, L. Peliti, Energy flow, partial equilibration, and effective temperatures in systems with slow dynamics, Physical Review E 55 (1997) 3898–3914.

[37] Y. Luo, S. Priyadarshini, M. Hsu, G. A. Worrell, J. Straub, D. Hsu, Velocity-dependent friction in a simple two-oscillator system. II. Regions of stability, companion paper, submitted (2026).

[38] B. Phipson, G. K. Smyth, Permutation p-values should never be zero: Calculating exact p-values when permutations are randomly drawn, Statistical Applications in Genetics and Molecular Biology 9 (1) (2010) Article 39. doi:10.2202/1544-6115.1585.

[39] S. Holm, A simple sequentially rejective multiple test procedure, Scandinavian Journal of Statistics 6 (2) (1979) 65–70. doi:10.2307/4615733.

[40] E. B. Wilson, Probable inference, the law of succession, and statistical inference, Journal of the American Statistical Association 22 (158) (1927) 209–212. doi:10.1080/01621459.1927.10502953.

[41] S. R. Cole, B. Voytek, Brain oscillations and the importance of waveform shape, Trends in Cognitive Sciences 21 (2) (2017) 137–149. doi:10.1016/j.tics.2016.12.008.

[42] L. L. Colgin, T. Denninger, M. Fyhn, T. Hafting, T. Bonnevie, O. Jensen, M.-B. Moser, E. I. Moser, Frequency of gamma oscillations routes flow of information in the hippocampus, Nature 462 (7271) (2009) 353–357. doi:10.1038/nature08573.

[43] M. A. Belluscio, K. Mizuseki, R. Schmidt, R. Kempter, G. Buzsáki, Cross-frequency phase-phase coupling between theta and gamma oscillations in the hippocampus, Journal of Neuroscience 32 (2) (2012) 423–435. doi:10.1523/JNEUROSCI.4122-11.2012.

[44] M. A. Kramer, U. T. Eden, Assessment of cross-frequency coupling with confidence using generalized linear models, Journal of Neuroscience Methods 220 (1) (2013) 64–74. doi:10.1016/j.jneumeth.2013.08.006.

[45] J. K. Nadalin, L.-E. Martinet, E. B. Blackwood, M.-C. Lo, A. S. Widge, S. S. Cash, U. T. Eden, M. A. Kramer, A statistical framework to assess cross-frequency coupling while accounting for confounding analysis effects, eLife 8 (2019) e44287. doi:10.7554/eLife.44287.

[46] T. Dupré la Tour, L. Tallot, L. Grabot, V. Doyère, V. van Wassenhove, Y. Grenier, A. Gramfort, Non-linear auto-regressive models for cross-frequency coupling in neural time series, PLoS Computational Biology 13 (12) (2017) e1005893. doi:10.1371/journal.pcbi.1005893.

[47] M. A. Kramer, A. B. L. Tort, N. J. Kopell, Sharp edge artifacts and spurious coupling in EEG frequency comodulation measures, Journal of Neuroscience Methods 170 (2) (2008) 352–357. doi:10.1016/j.jneumeth.2008.01.020.

[48] D. Dvorak, A. A. Fenton, Toward a proper estimation of phase-amplitude coupling in neural oscillations, Journal of Neuroscience Methods 225 (2014) 42–56. doi:10.1016/j.jneumeth.2014.01.002.

[49] D. Lozano-Soldevilla, N. ter Huurne, R. Oostenveld, Neuronal oscillations with non-sinusoidal morphology produce spurious phase-to-amplitude coupling and directionality, Frontiers in Computational Neuroscience 10 (2016) 87. doi:10.3389/fncom.2016.00087.

[50] H. Lei, N. A. Baker, X. Li, Data-driven parameterization of the generalized langevin equation, Proceedings of the National Academy of Sciences of the United States of America 113 (50) (2016) 14183–14188. doi:10.1073/pnas.1609587113.

[51] K. V. Mardia, P. E. Jupp, Directional Statistics, Wiley, Chichester, 2000.

