## Supplementary math for "Velocity-dependent friction in a simple two-oscillator system. I. Phase–amplitude coupling"

This document collects derivations and tests that support, but are not needed to follow, the main text. Equation, section and appendix numbers without the prefix S refer to the main text.

#### S1 Numerical scheme

**Update** The theta velocity is advanced with the semi-implicit trapezoidal step for its linear terms. The gamma coupling is integrated together with its baseline friction in one trapezoidal step, in which the friction acts on the average of the old and the new velocity; solving for the new velocity gives

$$\begin{aligned} v_{\gamma,n+1} &= \frac{(1-\frac{h}{2}c_n)v_{\gamma,n} - h\omega_\gamma^2 x_{\gamma,n} + \sigma\sqrt{h}\,\xi_{\gamma,n}}{1+\frac{h}{2}c_n}, \\ c_n &= g_{\gamma,0} - \frac{g_{vv}}{2}\big(v_{\theta,n}+v_{\theta,n+1}\big), \end{aligned} \tag{S1}$$

followed by $x_{n+1} = x_n + hv_{n+1}$ for both oscillators, with $h = 0.5\,\mathrm{ms}$ and independent standard normal $\xi$ [1].

**Contraction of gamma disturbances in continuous time** Along a fixed theta path, a small gamma disturbance $\delta u = (\delta x_\gamma, \delta v_\gamma)$ obeys the gamma equation without noise, $\delta\dot{u} = J(t)\,\delta u$ with $J = \big(\begin{smallmatrix} 0 & 1 \\ -\omega_\gamma^2 & -g_{\gamma,\mathrm{eff}}(t)\end{smallmatrix}\big)$. Liouville's formula gives the determinant of its propagator $\Phi(t)$, the factor by which a small area of gamma states is multiplied, from the trace of $J$:

$$\det\Phi(t) = \exp\Big(-\int_0^t g_{\gamma,\mathrm{eff}}\,\mathrm{d}s\Big) = \exp\big[-g_{\gamma,0}t + g_{vv}\big(x_\theta(t) - x_\theta(0)\big)\big].$$

The long-time contraction rate is the limit

$$\lim_{t\to\infty}\frac{1}{t}\ln\det\Phi(t) = -g_{\gamma,0} + g_{vv}\lim_{t\to\infty}\frac{x_\theta(t)-x_\theta(0)}{t} = -g_{\gamma,0},$$

because theta coordinate is stationary with finite variance $\sigma_x^2$ (Section S3), so that $x_\theta(t)/t \to 0$. This limit is the sum of the two Lyapunov exponents of gamma (the long-time growth rates of small disturbances), so $\lambda_1 + \lambda_2 = -g_{\gamma,0}$ for every coupling; Part II [2] discusses its consequences.

**The same property for the discrete step** One step of Eq. (S1) is the velocity update, which maps $(x_n, v_n)$ to $(x_n, v_{n+1})$ with determinant $a_n = (1-hc_n/2)/(1+hc_n/2)$, followed by the coordinate update, a shear with determinant one. Over $N = t/h$ steps, $\ln\prod_n a_n = \sum_n\big[-hc_n - (hc_n)^3/12 + \dots\big]$, and since $x_{n+1} = x_n + hv_{n+1}$,

$$h\sum_{n=0}^{N-1} c_n = g_{\gamma,0}t - g_{vv}\big[x_\theta(t) - x_\theta(0)\big] - \frac{g_{vv}h}{2}\big(v_{\theta,0} - v_{\theta,N}\big).$$

The last two terms stay bounded, so the discrete contraction rate is $-g_{\gamma,0} - h^2\langle c^3\rangle/12$, with $\langle c^3\rangle = g_{\gamma,0}^3 + 3g_{\gamma,0}g_{vv}^2\Theta_\theta$ for Gaussian theta velocities. At C1 the correction is $6\times10^{-4}\,\mathrm{s}^{-1}$, 0.06% of $g_{\gamma,0}$, whereas an explicit Euler step for the friction, with determinant $1-hc_n$, would shift the rate by $h\langle c^2\rangle/2 \approx 2.5\,\mathrm{s}^{-1}$, more than twice $g_{\gamma,0}$ itself. The scheme thus keeps $\lambda_1+\lambda_2 = -g_{\gamma,0}$, which Part II uses to measure the largest Lyapunov exponent without a numerical bias.

**Discrete check** At $g_{vv}=0$ the step is linear in $u_n=(x_n,v_n)$, $u_{n+1}=Au_n+B\xi_n$, with $\xi_n$ independent of $u_n$. Taking the covariance of both sides gives $\Sigma_{n+1}=A\Sigma_nA^\mathsf{T}+BB^\mathsf{T}$, whose fixed point $\Sigma = A\Sigma A^\mathsf{T}+BB^\mathsf{T}$ is the exact stationary covariance of the discrete map. For an oscillator of frequency $\omega$ and friction $g$ advanced by Eq. (S1), solving this $2\times2$ linear system gives

$$\Sigma_{vv}=\omega^2\Sigma_{xx}=\frac{\Theta}{1-(h\omega)^2/4},\qquad \Theta=\frac{\sigma^2}{2g},$$

so the step reproduces the continuous-time temperature up to the factor $1/[1-(h\omega)^2/4]$. At $\zeta=0.0027$ this gives $\Sigma_{vv}=147.37$ for theta, 0.004% above $\Theta_\theta$, and 9.912 for gamma, 0.9% above $\Theta_{\gamma,0}=\sigma^2/(2g_{\gamma,0})=9.824$.

### S2 The second-order equation of motion and its ten terms

This section derives the ten quadratic terms of Section 2.2 and shows which of them change the gamma amplitude.

#### S2.1 Expansion of stiffness and friction

Let $X(t)\in\mathbb{R}^N$ collect the collective coordinates, here the theta and gamma components, and $V=\dot X$ their velocities. Starting from the linear GLE $\ddot X = -KX - GV - \int_0^t H(\tau)V(t-\tau)\,\mathrm{d}\tau + F(t)$ [3], we let the stiffness and the friction depend on the state and keep the first order in $X$ and $V$:

$$\begin{aligned}\ddot X &= -K(X,\dot X)\,X - G(X,\dot X)\,\dot X - \int_0^t H(\tau)\,\dot X(t-\tau)\,\mathrm{d}\tau + F(t),\\ K(X,\dot X)_{ij} &= K_{0,ij} + \textstyle\sum_k \big(K_{xx,ijk}X_k + K_{vx,ijk}\dot X_k\big),\\ G(X,\dot X)_{ij} &= G_{0,ij} + \textstyle\sum_k \big(G_{xv,ijk}X_k + G_{vv,ijk}\dot X_k\big).\end{aligned}\tag{S2}$$

The right-hand side then contains every product of two of the coordinates and velocities: $XX$ with $K_{xx}$, $VV$ with $G_{vv}$, and $XV$ with $K_{vx}$ or $G_{xv}$. Only the sum of the two coordinate–velocity coefficients for a given product enters the equation, so they can be collected in one of them for estimation. For two oscillators there are ten distinct products, the terms of Section 2.2. Without memory, the power balance of Section S3, applied to every component, fixes the noise, $\langle F(t)F(0)^\mathsf{T}\rangle = 2G\langle VV^\mathsf{T}\rangle\delta(t)$ for a symmetric friction. The stiffness follows from stationarity: $\mathrm{d}\langle VX^\mathsf{T}\rangle/\mathrm{d}t = \langle \ddot XX^\mathsf{T}\rangle + \langle VV^\mathsf{T}\rangle = 0$, and multiplying the equation of motion by $X^\mathsf{T}$ and averaging gives $\langle\ddot XX^\mathsf{T}\rangle = -K\langle XX^\mathsf{T}\rangle$ when $\langle VX^\mathsf{T}\rangle$, $\langle FX^\mathsf{T}\rangle$ and the third moments of $X$ vanish, so that $K=\langle VV^\mathsf{T}\rangle\langle XX^\mathsf{T}\rangle^{-1}$; for one oscillator these read $\sigma^2=2G\langle v^2\rangle$ and $\omega^2=\langle v^2\rangle/\langle x^2\rangle$. Because every coefficient multiplies a known function of the data, all of them can be estimated by linear regression.

#### S2.2 What each quadratic term does to gamma

To see which terms change the gamma amplitude, we use the amplitude–phase variables of Eq. (A.1), in which $r^2/2$ is the gamma energy and $\sin^2\varphi$ its kinetic share. For a gamma equation $\dot v_\gamma = -\omega_\gamma^2 x_\gamma + f$ with an additional force $f$, differentiating Eq. (A.1) and solving for $\dot r$ and $\dot\varphi$ gives

$$\dot r = -f\sin\varphi,\qquad \dot\varphi = \omega_\gamma - \frac{f}{r}\cos\varphi.\tag{S3}$$

Each quadratic term of the gamma equation is such a force. Averaging over $\varphi$ at fixed theta state [4]:

- for $f = c(\theta)v_\gamma = -c(\theta)r\sin\varphi$, $\overline{\dot r/r} = c(\theta)\overline{\sin^2\varphi} = c(\theta)/2$, so the friction-type terms with $c(\theta) = g_{vv}v_\theta + g_{xv}x_\theta$ modulate the amplitude;
- for $f = -k(\theta)x_\gamma$, $\bar{\dot r} = k(\theta)\omega_\gamma^{-1}\overline{\cos\varphi\sin\varphi}\, r = 0$ and $\bar{\dot\varphi} = \omega_\gamma + k(\theta)/(2\omega_\gamma)$, so the stiffness-type terms with $k(\theta) = K_{xx}x_\theta + K_{vx}v_\theta$ modulate the frequency at fixed energy;
- for theta-only terms, $f$ contains only the frequencies 0, $\omega_\theta$ and $2\omega_\theta$, far from $\omega_\gamma$, so $\bar{\dot r} = 0$;
- for gamma-only terms, $\bar{\dot r}$ is proportional to averages of $\sin^3\varphi$, $\cos\varphi\sin^2\varphi$ or $\cos^2\varphi\sin\varphi$, which vanish.

Linear cross terms are of the theta-only kind in the gamma equation and give no first-order effect either. Stiffness-type terms add no energy at first order (second item above). At the next order, a slow change of the gamma frequency conserves the action $E_\gamma/\omega$, an adiabatic invariant; with $E_\gamma = \frac{1}{2}\omega^2 A^2$ the coordinate amplitude then follows $A \propto \omega_{\text{eff}}^{-1/2}$, so that $\ln A$ changes by $-\Delta\omega/(2\omega_\gamma)$, which does not accumulate over the theta cycle. This is a modulation of the form treated in Appendix A.4, with depth $\kappa_s = \Delta\omega_{\max}/(2\omega_\gamma)$, and its small-$\kappa$ limit gives $\mathrm{MI} \approx \kappa_s^2/(4\ln N_\mathrm{b})$, second order in the frequency-modulation depth; the preferred phase is the theta phase of lowest gamma frequency. For the swing of the gamma frequency by 17 Hz reported in Section 5.6, $\kappa_s \approx 0.07$ and $\mathrm{MI} \approx 4\times10^{-4}$, as measured. In the theta equation, averaging over the gamma phase leaves only terms with two gamma factors, which force theta in proportion to gamma energy (back-reaction), while theta's own quadratic terms do not average out and make theta non-sinusoidal.

##### S2.3 The ten-term test

Table S1 lists the results of the ten-term test of Appendix B.

Table S1: **The ten second-order terms at equal force strength** ($\zeta = 0.0027$, seeds 100–149).

| Term | Class | $c$ | Complete | Median MI [IQR] | Phase (rad) | $R$ |
|---|---|---|---|---|---|---|
| $v_\theta v_\gamma$ | friction | 8.20 | 50/50 | 0.213 [0.139, 0.285] | −0.014 | 1.0000 |
| $x_\theta v_\gamma$ | friction | 206.1 | 50/50 | 0.212 [0.141, 0.284] | 1.553 | 1.0000 |
| $x_\theta x_\gamma$ | stiffness | $7.77\times10^4$ | 50/50 | $4.0\times10^{-4}$ [2.2, 8.8] | 0.095 | 0.73 |
| $v_\theta x_\gamma$ | stiffness | $3.09\times10^3$ | 50/50 | $4.3\times10^{-4}$ [1.9, 9.0] | −1.504 | 0.70 |
| $x_\theta^2$ | single | 772 | 50/50 | $6.2\times10^{-5}$ [4.1, 9.2] | 0.826 | 0.09 |
| $x_\theta v_\theta$ | single | 53.2 | 50/50 | $6.2\times10^{-5}$ [4.0, 8.9] | 1.263 | 0.07 |
| $v_\theta^2$ | single | 1.22 | 50/50 | $5.8\times10^{-5}$ [4.4, 9.2] | 0.507 | 0.06 |
| $x_\gamma^2$ | single | $2.61\times10^6$ | 36/50 | $9.2\times10^{-5}$ [4.2, 12.4] | 0.781 | 0.10 |
| $x_\gamma v_\gamma$ | single | $1.20\times10^4$ | 50/50 | $5.5\times10^{-5}$ [3.2, 8.9] | −2.241 | 0.13 |
| $v_\gamma^2$ | single | 18.3 | 50/50 | $9.7\times10^{-5}$ [6.1, 16.0] | 2.374 | 0.17 |

The term enters the gamma equation as $+cm$, with $c$ set to equal force RMS (Section 4.3). Interquartile ranges of small MI values are given in units of the stated power of ten. Phase: circular mean of per-record preferred phases (0 = theta coordinate peak); $R$: their mean resultant length (consistency). Single: single-oscillator term. Fourteen $x_\gamma^2$ records escaped over the finite barrier of the cubic potential that this term generates.

#### S3 Theta: temperature, response and amplitude distribution

This section proves Eq. (4) and the Rayleigh law of theta amplitudes used in Appendix A.4; Table S2 lists the symbols of the derivations in Appendix A of the main text and in Sections S2 to S5.

Table S2: **Symbols used in the derivations.**

| Symbol | Meaning | Defined in |
|---|---|---|
| $E_\theta$, $E_\gamma$ | oscillator energies, $\frac{1}{2}(v^2+\omega^2x^2)$ | Section S3, Appendix A.1 |
| $\Theta_\theta$, $\sigma_x$ | theta effective temperature $\langle v_\theta^2\rangle$; typical theta amplitude $\sqrt{\Theta_\theta}/\omega_\theta$ | Section S3 |
| $C_\theta$, $\rho_\theta$, $\chi_\theta$ | theta coordinate correlation $\langle x_\theta(t)x_\theta(0)\rangle$, its normalized form $C_\theta/C_\theta(0)$, and the response of $x_\theta$ to a unit impulse | Section S3 |
| $a_\theta$, $D_a$ | theta (Hilbert) amplitude $(x_\theta^2+v_\theta^2/\omega_\theta^2)^{1/2}$; its diffusion coefficient $\sigma^2/(2\omega_\theta^2)$ | Section S3 |
| $r$, $\varphi$ | gamma amplitude in velocity units (the $a_\gamma$ of the main text) ($r^2/2=E_\gamma$) and fast gamma phase | Appendix A.1 |
| $\phi_\theta$ | theta phase, zero at the theta coordinate peak | Appendix A.4 |
| $\kappa_a$, $\kappa$ | PAC parameter of one theta cycle, $g_{vv}a_\theta/2$; of the parameters, $g_{vv}\sqrt{2\Theta_\theta}/(2\omega_\theta)$ | Appendix A.4 |
| $N_\mathrm{b}$ | number of theta-phase bins of the modulation index (18) | Appendix A.4, Section S6 |
| $\Theta_{\gamma,0}$ | gamma temperature without coupling, $\sigma^2/(2g_{\gamma,0})$ | Section S5 |
| $\tau$, $\Delta_\tau$, $\nu$ | time lag; theta displacement $x_\theta(t)-x_\theta(t-\tau)$; $\nu=g_{vv}^2\sigma_x^2$ | Section S5 |
| $\eta_\mathrm{FD}$, $\alpha_E$, $F$ | effective gamma friction; energy-weighted theta velocity; variance factor $\langle v_\gamma^2\rangle/\Theta_{\gamma,0}$ | Section S5 |

**Temperature** The theta oscillator, $\mathrm{d}x_\theta = v_\theta\,\mathrm{d}t$ and $\mathrm{d}v_\theta = (-\omega_\theta^2 x_\theta - g_{\theta,0}v_\theta)\,\mathrm{d}t + \sigma\,\mathrm{d}W_\theta$, is a two-dimensional Ornstein–Uhlenbeck process. Itô's formula (the chain rule for noise-driven variables) for its energy $E_\theta = \frac{1}{2}(v_\theta^2+\omega_\theta^2x_\theta^2)$ gives

$$\mathrm{d}E_\theta = \Big(-g_{\theta,0}v_\theta^2 + \frac{\sigma^2}{2}\Big)\mathrm{d}t + \sigma v_\theta\,\mathrm{d}W_\theta.$$

In the stationary state the mean energy is constant and the stochastic term has zero mean, so the power $\sigma^2/2$ injected by the noise equals the power $g_{\theta,0}\langle v_\theta^2\rangle$ removed by friction. This balance alone fixes $\langle v_\theta^2\rangle$. The coordinate variance follows from the other two quadratic moments. Because $x_\theta$ itself carries no noise, Itô's formula gives $\mathrm{d}(x_\theta^2) = 2x_\theta v_\theta\,\mathrm{d}t$ and $\mathrm{d}(x_\theta v_\theta) = (v_\theta^2 - \omega_\theta^2 x_\theta^2 - g_{\theta,0}x_\theta v_\theta)\,\mathrm{d}t + \sigma x_\theta\,\mathrm{d}W_\theta$, so that in the stationary state

$$0 = \frac{\mathrm{d}\langle x_\theta^2\rangle}{\mathrm{d}t} = 2\langle x_\theta v_\theta\rangle, \qquad 0 = \frac{\mathrm{d}\langle x_\theta v_\theta\rangle}{\mathrm{d}t} = \langle v_\theta^2\rangle - \omega_\theta^2\langle x_\theta^2\rangle - g_{\theta,0}\langle x_\theta v_\theta\rangle.$$

The first equation states that coordinate and velocity are uncorrelated: in the stationary state theta is as likely to move outward as inward. The second then gives $\langle v_\theta^2\rangle = \omega_\theta^2\langle x_\theta^2\rangle$, the equipartition of the mean kinetic and potential energies, so that the standard deviation of theta coordinate, $\sigma_x \equiv \langle x_\theta^2\rangle^{1/2}$, follows from the temperature:

$$\boxed{\Theta_\theta \equiv \langle v_\theta^2\rangle = \frac{\sigma^2}{2g_{\theta,0}}, \qquad \sigma_x = \frac{\sqrt{\Theta_\theta}}{\omega_\theta}.} \tag{S4}$$

Written as $\sigma^2 = 2g_{\theta,0}\Theta_\theta$, Eq. (S4) is the Einstein relation between noise and friction, with the theta effective temperature $\Theta_\theta$ in place of $k_BT/m$ [5–7]. In the model $\sigma$ and $g_{\theta,0}$ are given, and $\Theta_\theta$ is the temperature that makes them consistent; at $\zeta = 0.0027$, $\Theta_\theta = 147.4$ and $\sigma_x = 0.483$.

**Response and correlation** The same friction that limits the spontaneous fluctuations also damps the response to a push, so the temperature fixes both. Both the response of $x_\theta$ to a unit impulse of force at $t=0$, $\chi_\theta$, and the stationary correlation $C_\theta(t) = \langle x_\theta(t)x_\theta(0)\rangle$ obey the free damped-oscillator equation $\ddot{y} + g_{\theta,0}\dot{y} + \omega_\theta^2 y = 0$ for $t>0$; for $C_\theta$ this holds because future noise is uncorrelated with $x_\theta(0)$. They differ only in their initial values. The impulse changes the velocity by one but not the coordinate, so $\chi_\theta(0)=0$ and $\dot{\chi}_\theta(0)=1$, whereas the moments above give $C_\theta(0) = \langle x_\theta^2 \rangle = \Theta_\theta/\omega_\theta^2$ and $C'_\theta(0) = \langle x_\theta v_\theta \rangle = 0$. With $\Omega_d = (\omega_\theta^2 - g_{\theta,0}^2/4)^{1/2}$, solving the two initial-value problems gives, for $t \geq 0$,

$$\chi_\theta(t) = e^{-g_{\theta,0}t/2}\,\frac{\sin \Omega_d t}{\Omega_d}, \qquad C_\theta(t) = \frac{\Theta_\theta}{\omega_\theta^2}\, e^{-g_{\theta,0}t/2}\Big[\cos \Omega_d t + \frac{g_{\theta,0}}{2\Omega_d}\sin \Omega_d t\Big],$$

The derivative $C'_\theta$ obeys the same equation, with $C'_\theta(0) = 0$ and $C''_\theta(0) = -\omega_\theta^2 C_\theta(0) = -\Theta_\theta$, which are $-\Theta_\theta$ times the initial values of $\chi_\theta$; since the solution is fixed by its initial values,

$$\chi_\theta(t) = -\frac{1}{\Theta_\theta}\,\frac{\mathrm{d}C_\theta(t)}{\mathrm{d}t}, \qquad t \geq 0. \tag{S5}$$

Theta therefore responds to a brief push exactly as its spontaneous fluctuations decay, with $\Theta_\theta$ as the conversion factor: Eq. (S5) is the fluctuation–dissipation theorem with $\Theta_\theta$ in place of $k_B T/m$. Section 5.2 tests it.

**Amplitude distribution** A linear system driven by Gaussian noise has a Gaussian stationary state, and uncorrelated Gaussian variables are independent; by the moments above, $x_\theta$ and $y = v_\theta/\omega_\theta$ are therefore independent Gaussians with zero mean and variance $\sigma_x^2$, so the theta (Hilbert) amplitude $a_\theta = (x_\theta^2 + y^2)^{1/2}$ follows the Rayleigh distribution [8]

$$p(a_\theta) = \frac{a_\theta}{\sigma_x^2}\exp\Big(-\frac{a_\theta^2}{2\sigma_x^2}\Big),$$
$$\Pr(a_\theta > k\sigma_x) = e^{-k^2/2}, \qquad \text{median}\, a_\theta = \sigma_x\sqrt{2\ln 2}. \tag{S6}$$

Equivalently, the theta energy $E_\theta = \frac{1}{2}\omega_\theta^2 a_\theta^2$ is exponentially (Boltzmann) distributed with mean $\Theta_\theta$. Since $\langle x_\theta^2\rangle = \langle a_\theta^2\rangle/2$, $\sigma_x$ is the scale of this distribution, which is why we call it the typical theta amplitude; the root-mean-square amplitude is $\sqrt{2}\,\sigma_x$ and the median $\sigma_x\sqrt{2\ln 2}$. A cycle three times larger than typical ($k = 3$) occurs with probability 1.1%.

**Amplitude dynamics** Averaging the energy equation above over the theta phase replaces $v_\theta^2$ by $E_\theta$ in the drift and in the noise variance, $\mathrm{d}E_\theta = (-g_{\theta,0}E_\theta + \sigma^2/2)\,\mathrm{d}t + \sigma\sqrt{E_\theta}\,\mathrm{d}B$, and Itô's formula for $a_\theta = \sqrt{2E_\theta}/\omega_\theta$ then gives

$$\mathrm{d}a_\theta = \Big(-\frac{g_{\theta,0}}{2}a_\theta + \frac{D_a}{2a_\theta}\Big)\mathrm{d}t + \sqrt{D_a}\,\mathrm{d}B, \qquad D_a = \frac{\sigma^2}{2\omega_\theta^2},$$

The stationary solution of the corresponding equation for the probability density (the Fokker–Planck equation) is $p \propto a_\theta \exp[-g_{\theta,0}a_\theta^2/(2D_a)]$, which is Eq. (S6) because $D_a/g_{\theta,0} = \sigma_x^2$, and the linear drift $-g_{\theta,0}a_\theta/2$ sets the relaxation time $2/g_{\theta,0} = 29.5\,\mathrm{s}$ at $\zeta = 0.0027$. A 10 s record therefore samples essentially one theta amplitude.

### S4 Both friction-type terms

Steps 2–4 of Appendix A extend directly to a friction that follows both theta velocity and theta coordinate. With both friction-type terms, $g_{\gamma,\mathrm{eff}} = g_{\gamma,0} - g_{vv}v_\theta - g_{xv}x_\theta$; the averaged rule $\mathrm{d}\ln r/\mathrm{d}t = -g_{\gamma,\mathrm{eff}}/2$ of Appendix A.2 gives the first line below, and integrating the theta equation of motion over time expresses $\int x_\theta\,\mathrm{d}s$ through the changes of theta velocity and coordinate (second line):

$$\ln\frac{r(t)}{r(0)} = -\frac{g_{\gamma,0}t}{2} + \frac{g_{vv}}{2}\Delta x_\theta + \frac{g_{xv}}{2}\int_0^t x_\theta\,\mathrm{d}s,$$
$$\int_0^t x_\theta\,\mathrm{d}s = -\frac{\Delta v_\theta + g_{\theta,0}\Delta x_\theta - \sigma\Delta W_\theta}{\omega_\theta^2}. \tag{S7}$$

Within a theta cycle, with $x_\theta = a\cos\phi$, $v_\theta = -a\omega_\theta \sin\phi$ and the term $g_{\theta,0}\Delta x_\theta$, smaller than $\Delta v_\theta$ by the factor $g_{\theta,0}/\omega_\theta = \zeta$, neglected, the exponent is a cosine plus a sine of the theta phase, which combine into one shifted cosine:

$$A_\gamma(\phi) \propto \exp\left[\frac{a}{2}\Big(g_{vv}\cos\phi + \frac{g_{xv}}{\omega_\theta}\sin\phi\Big)\right] = e^{\kappa_a \cos(\phi-\delta)},$$
$$g_{\mathrm{c}} = \sqrt{g_{vv}^2 + \Big(\frac{g_{xv}}{\omega_\theta}\Big)^2}, \qquad \delta = \operatorname{atan2}\Big(\frac{g_{xv}}{\omega_\theta}, g_{vv}\Big), \tag{S8}$$

with $\kappa_a = a g_{\mathrm{c}}/2$. The modulation index does not depend on $\delta$, so Eqs. (A.7) and (6) hold with $g_{vv} \to g_{\mathrm{c}}$. The noise term adds a slow multiplicative factor when $g_{xv} \neq 0$, which relaxes on the time scale $1/g_{\gamma,0}$ and is largely absorbed by the log-linear detrending. Table S3 compares the generalized closed form with simulation.

Table S3: **Generalized closed form against simulation** ($\zeta = 0.0027$, 100 records each).

| Case | $g_{vv}$ | $g_{xv}$ | Median MI | Predicted | Phase (rad) | Predicted |
|---|---|---|---|---|---|---|
| $v_\theta v_\gamma$ | 8.2 | 0 | 0.222 | 0.233 | −0.01 | 0 |
| $x_\theta v_\gamma$ | 0 | $8.2\,\omega_\theta$ | 0.217 | 0.233 | 1.55 | 1.571 |
| Mixed, 45° | $8.2/\sqrt{2}$ | $8.2\,\omega_\theta/\sqrt{2}$ | 0.219 | 0.233 | 0.77 | 0.785 |
| Both, $g_{\mathrm{c}} = 8.2\sqrt{2}$ | 8.2 | $8.2\,\omega_\theta$ | 0.301 | 0.314 | 0.78 | 0.785 |

#### S5 Gamma fluctuation–dissipation relation and departure from equilibrium

This section derives the gamma variance of Fig. 3 and defines the response temperature of Table S4. Taking the full expectation of the gamma energy equation of Appendix A.3 in the stationary state gives an exact power balance, which has the form of the theta relation of Section S3 with an effective friction $\eta_{\mathrm{FD}}$:

$$0 = -g_{\gamma,0}\langle v_\gamma^2\rangle + g_{vv}\langle v_\theta v_\gamma^2\rangle + \frac{\sigma^2}{2},$$
$$\langle v_\gamma^2\rangle = \frac{\sigma^2}{2\eta_{\mathrm{FD}}}, \qquad \eta_{\mathrm{FD}} = g_{\gamma,0} - g_{vv}\alpha_E, \qquad \alpha_E = \frac{\langle v_\theta v_\gamma^2\rangle}{\langle v_\gamma^2\rangle}. \tag{S9}$$

The three terms of the balance are the power removed by the baseline friction, the power pumped into gamma by the theta-modulated friction, and the power injected by the noise; $\alpha_E$ is the theta velocity weighted by the gamma kinetic energy. Equation (S9) is exact whenever the second moment exists, and $\eta_{\mathrm{FD}} > 0$ whenever it does. It is not closed, however, because $\alpha_E$ is unknown. Gamma energy grows with the recent theta displacement, which is positively correlated with the present theta velocity, so $\alpha_E > 0$ and the effective friction is lowered.

To evaluate it, we average Eq. (A.5), which gives the gamma energy along any theta path, over the Gaussian theta paths, use $\langle v_\gamma^2\rangle \approx \langle E_\gamma\rangle$ and write $\tau = t - s$ for the time lag. The exponent then contains the theta displacement $\Delta_\tau = x_\theta(t) - x_\theta(t-\tau)$, which is Gaussian with zero mean and variance $\langle \Delta_\tau^2\rangle = 2\langle x_\theta^2\rangle - 2C_\theta(\tau) = 2\sigma_x^2[1-\rho_\theta(\tau)]$; the Gaussian identity $\mathbb{E}[e^Z] = e^{\langle Z^2\rangle/2}$ for a zero-mean Gaussian $Z$ then gives $\mathbb{E}[e^{g_{vv}\Delta_\tau}] = e^{\nu[1-\rho_\theta(\tau)]}$ with $\nu = g_{vv}^2\sigma_x^2$. Relative to the gamma temperature without coupling, $\Theta_{\gamma,0} = \sigma^2/(2g_{\gamma,0})$,

$$\frac{\langle v_\gamma^2\rangle}{\Theta_{\gamma,0}} = \frac{g_{\gamma,0}}{\eta_{\mathrm{FD}}} = F(g_{vv}) = g_{\gamma,0}\int_0^\infty e^{-g_{\gamma,0}\tau} e^{\nu[1-\rho_\theta(\tau)]}\mathrm{d}\tau \approx e^\nu I_0(\nu). \tag{S10}$$

The last step uses $\rho_\theta(\tau) \approx \cos\omega_\theta\tau$ for $\tau \lesssim 1/g_{\gamma,0}$ and, because $e^{-g_{\gamma,0}\tau}$ varies slowly over a theta period, the phase average $(2\pi)^{-1}\int_0^{2\pi} e^{\nu(1-\cos\psi)}\mathrm{d}\psi = e^\nu I_0(\nu)$. Since $0 \le 1-\rho_\theta \le 2$,

$g_{\gamma,0}e^{-2\nu} \le \eta_{\rm FD} \le g_{\gamma,0}$: the lower bound vanishes only as $\nu \to \infty$, so within this approximation the coupling can make the effective friction arbitrarily small but never zero, whereas a constant $\alpha_E = \mathrm{RMS}(v_\theta)$ would give $\eta_{\rm FD} = 0$ already at $g_{vv} = g_{\gamma,0}/\sqrt{\Theta_\theta} = 0.084$. For large $\nu$ the mean is dominated by rare large $\Delta_\tau$, and the expression assumes that gamma keeps oscillating in those cycles (Appendix A.2). Strictly, the second moment diverges near $g_{vv} \approx 0.4$, where $\eta_{\rm FD} \to 0$, but only because of theta amplitudes of negligible probability [2]; Eq. (S10) describes the bulk of the distribution, which finite samples measure (Table S4).

**Departure from equilibrium** The variance alone cannot show whether gamma is in equilibrium, because an equilibrium oscillator with friction $\eta_{\rm FD}$ would have the same variance. The response can. In equilibrium the fluctuation–dissipation theorem fixes the ratio of the spontaneous fluctuations at a given frequency to the dissipative part of the response at that frequency; this ratio, normalized so that it equals the kinetic temperature in equilibrium, defines a response temperature $\Theta_{\rm resp}$ [9]. We evaluate it for gamma at 57–63 Hz, with the response to a brief push propagated along the same theta paths (Section 4.3). Without coupling $\Theta_{\rm resp}$ equals the kinetic temperature; with coupling it falls increasingly below it (Table S4), because the energy pumped in by the theta-modulated friction raises the fluctuations without a matching increase of the response.

Table S4 lists the values shown in Fig. 3. At $g_{vv} = 4$ the confidence interval is wide because rare large excursions slow the convergence of the sample mean, but it contains the prediction.

Table S4: **Gamma fluctuation–dissipation relation** ($\zeta = 0.0027$).

| $g_{vv}$ | $\langle v_\gamma^2\rangle/\Theta_{\gamma,0}$ [95% CI] | $F(g_{vv})$, Eq. (S10) | $\eta_{\rm FD}/g_{\gamma,0} = 1/F$ | $\Theta_{\rm resp}/\Theta_{\gamma,0}$, 57–63 Hz |
|---|---|---|---|---|
| 0 | 0.997 [0.962, 1.026] | 1 | 1 | 0.985 |
| 0.5 | 1.076 [1.039, 1.112] | 1.061 | 0.943 | 1.017 |
| 1 | 1.350 [1.277, 1.432] | 1.278 | 0.782 | 1.144 |
| 2 | 3.64 [3.07, 4.29] | 3.08 | 0.324 | 1.95 |
| 3 | 21.4 [14.4, 30.3] | 19.0 | 0.053 | 4.99 |
| 4 | 234 [113, 391] | 340 | 0.0029 | 25.9 |

Seeds 4101–4120, 200 s after burn-in; intervals are bootstrap percentiles over seeds. $\Theta_{\gamma,0} = 9.824$.

#### S6 PAC estimator

For a record of $N$ samples at $t_n = nh$, the centred coordinates $\tilde{x}_i = x_i - \bar{x}_i$ give analytic signals $z_i = \tilde{x}_i + \mathrm{i}\mathcal{H}\{\tilde{x}_i\}$, computed by FFT, with $\phi_\theta = \arg z_\theta$ and $A_\gamma = |z_\gamma|$. In the ten-term test, $\tilde{x}_\gamma$ is first band-passed by setting its FFT coefficients outside 40–80 Hz to zero. A log-linear trend $\ln A_\gamma = \beta_0 + \beta_1 t + \ln U$ is fitted by least squares over samples with $A_\gamma > 10^{-3} \max A_\gamma$, and the detrended envelope is $A_\gamma / \exp(\hat\beta_0 + \hat\beta_1 t)$. Theta phase is divided into $N_{\rm b} = 18$ equal bins $B_k$ with centres $\psi_k$; with $\bar{A}_k$ the mean envelope in bin $k$ and $p_k = \bar{A}_k / \sum_j \bar{A}_j$,

$$\mathrm{MI} = \frac{1}{\ln N_{\rm b}}\Big(\ln N_{\rm b} + \sum_{k=1}^{N_{\rm b}} p_k \ln p_k\Big) = \frac{D_{\rm KL}(\boldsymbol{p}\,\|\,\boldsymbol{u})}{\ln N_{\rm b}}, \qquad \phi_{\rm pref} = \arg \sum_{k=1}^{N_{\rm b}} \bar{A}_k\, e^{\mathrm{i}\psi_k}, \tag{S11}$$

and a record is invalid if any bin is empty. Surrogates identify theta cycles from phase wraps and circularly shift the envelope within each cycle by an independent random offset, applying the same offsets to raw and detrended envelopes.

#### References


[1] P. E. Kloeden, E. Platen, Numerical Solution of Stochastic Differential Equations, Springer, Berlin, 1992.

[2] Y. Luo, S. Priyadarshini, M. Hsu, G. A. Worrell, J. Straub, D. Hsu, Velocity-dependent friction in a simple two-oscillator system. II. Regions of stability, companion paper, submitted (2026).

[3] D. Hsu, M. Hsu, Zwanzig–Mori projection operators and EEG dynamics: Deriving a simple equation of motion, PMC Biophysics 2 (2009) 6. doi:10.1186/1757-5036-2-6.

[4] J. A. Sanders, F. Verhulst, J. Murdock, Averaging Methods in Nonlinear Dynamical Systems, 2nd Edition, Vol. 59 of Applied Mathematical Sciences, Springer, New York, 2007.

[5] R. Kubo, The fluctuation-dissipation theorem, Reports on Progress in Physics 29 (1) (1966) 255–284. doi:10.1088/0034-4885/29/1/306.

[6] C. W. Gardiner, Handbook of Stochastic Methods for Physics, Chemistry and the Natural Sciences, 3rd Edition, Springer, Berlin, 2004.

[7] H. Risken, The Fokker–Planck Equation: Methods of Solution and Applications, 2nd Edition, Springer, Berlin, 1989.

[8] S. O. Rice, Mathematical analysis of random noise, Bell System Technical Journal 23 (3) (1944) 282–332.

[9] L. F. Cugliandolo, J. Kurchan, L. Peliti, Energy flow, partial equilibration, and effective temperatures in systems with slow dynamics, Physical Review E 55 (1997) 3898–3914.
